# A putative bacterial ecocline in *Klebsiella pneumoniae*

**DOI:** 10.1101/2025.01.20.633859

**Authors:** Siqi Liu, Sarah L. Svensson, Daniel Falush

**Affiliations:** Shanghai Institute of Materia Medica, Chinese Academy of Sciences, 320 Yueyang Road, Shanghai

## Abstract

The genetic structure of bacterial species is most often interpreted in terms of demographic processes such as clonal descent but can also reflect natural selection and hence give functional and ecological insight. *Klebsiella pneumoniae* (KP) disperses effectively around the world and has high recombination rates, which should result in the species having a well-mixed gene pool. Nevertheless, phylogenies based on diverse KP strains contain a "backbone". This structure reflects a component of variation where the first component in Principal Components Analysis (PCA), PC1, explains 16.8% of the total variation. We propose that the component reflects a “bacterial ecocline” generated by diversifying selection on a quantitative genetic trait. We simulated a model in which a trait is influenced by many genes, and strains with the most extreme trait values have a small advantage, which can recapitulate our KP PCA results and other features of its genetic diversity. As well as providing an explanation for the phylogenetic backbone, our results provide insight into how species such as KP can speciate, via stronger selection on the trait or a reduction in gene flow. Our hypothesis that there is a bacterial ecocline in KP raises two questions, namely what the trait is underlying it and why is the trait under diversifying selection? The genes that are most strongly associated with PC1 provide some hints, with the top locus encoding Kpa fimbriae. Identification of the trait, if it exists, should facilitate insight into selection on quantitative genetic traits in natural bacterial populations, which have largely been unstudied in microbiology, except in the atypical context of antibiotic resistance.

## Introduction

What information can be gleaned from phylogenetic trees[1]? The shape of a tree reflects key demographic factors, such as clonal relationships, geographic structure shaped by barriers to dispersal and genetic exchange[2]. Deep splits within trees and an absence of intermediate strains can for example reflect long-standing divergence between groups that has arisen due to niche specialization and/or low levels of genetic exchange[3,4]. However, many species show a pattern of differentiation that is more continuous in character, which can be harder to explain.

*Klebsiella pneumoniae* (KP) is a major human pathogen capable of causing severe infections in immunocompromised individuals[5], but is found in a wide variety of environments[6], including soil[7], wastewater[8], and even the ocean[9]. Like many other bacterial species, KP has an enormous population size, exhibits high transmission efficiency and has high recombination rates between strains[10]. As a result, its global genetic diversity can be captured from samples collected in a single region, whether from Europe[11], Africa[12], Asia[13], or elsewhere. Despite these features, as we will describe in detail below, the phylogenetic tree of a broad sample of KP strains shows deep structure, with strains radiating from multiple points along a "backbone". Until now, to our knowledge, no explanations have been proposed for this shape.

We propose that the shape of the phylogenetic tree of KP reflects diversifying selection[14] acting on a quantitative trait[15]. Specifically, we simulate a population of bacteria with a single phenotypic trait. Genetic variants influence the trait additively, meaning that specific variants increasing or decrease trait values by a fixed amount that does not depend on the presence or absence of other variants. We show that if strains exhibiting extreme trait values have a fitness advantage, we can recapitulate patterns of diversity found in KP. We call the gradient in variant frequency values that arises a "bacterial ecocline". Quantitative traits are central to our understanding of genetic diversity in animals and plants but have hitherto been largely overlooked for bacteria. If we can identify the trait or traits under selection in KP, it should provide considerable insight into the genetic architecture of bacterial quantitative traits as well as the fitness trade-offs[16] that the species experiences.

## Results

### The phylogenetic backbone reflects a component of variation

Genetic relationships amongst bacteria are most often represented using a phylogenetic tree[1,17]. Trees based on core genome SNPs from global collections of *Klebsiella pneumoniae* (KP) isolates are "star-like" (Figure 1a & Supplementary Figure 1, non-redundant dataset with clonally related strains removed; Supplementary Figure 2, entire dataset), with many branches of approximately equal length. This property of the tree reflects extensive recombination, with the length of the branches reflecting the overall level of diversity in the gene pool[10,18]. However, the overall shape of the KP tree is oval rather than approximately circular as for example seen in *Vibrio parahaemolyticus*[19,20] and KP lineages, instead of radiating from a single point, are connected by a "backbone" (Figure 1a & Supplementary Figure 2). This backbone is present in any phylogeny drawn based on a diverse sample of KP genomes (Figure 1a and Supplementary Figure 2) but it may be less visually evident than in Figure 1a, depending on the strains sampled, the rooting of the tree, and the phylogenetic and plotting algorithms.

**Figure 1.**
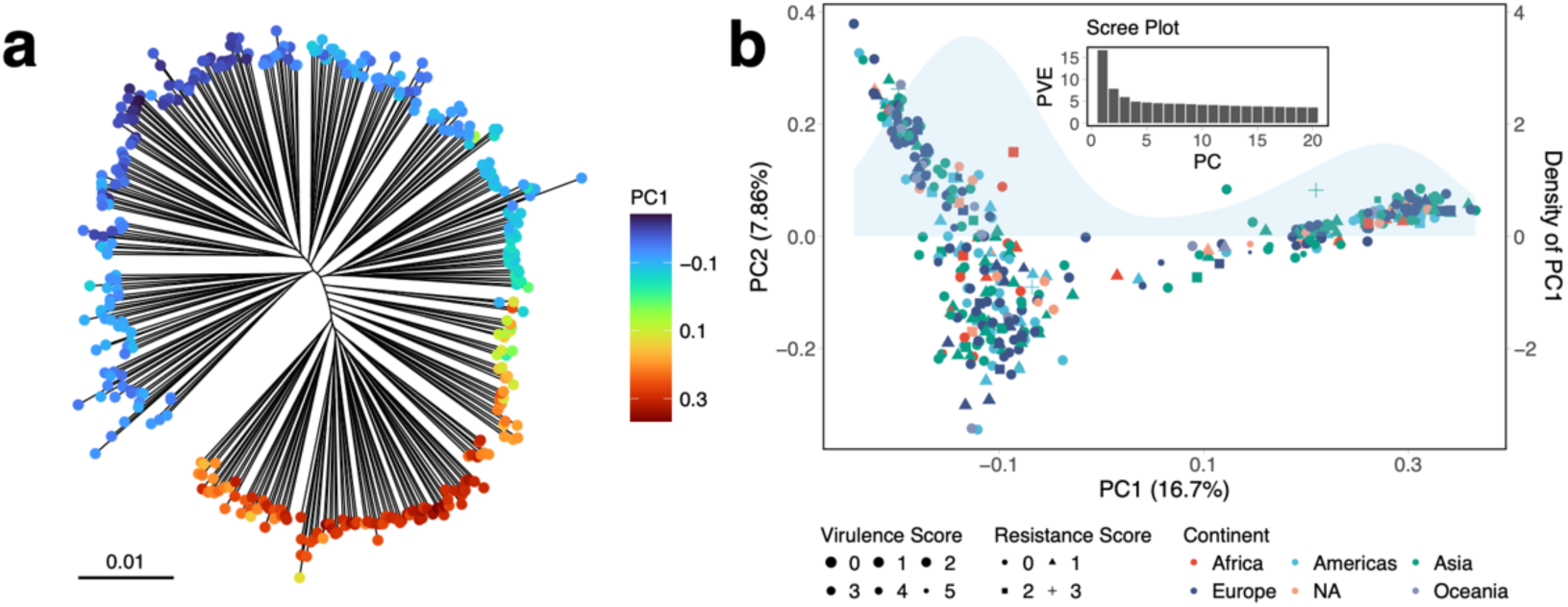
Population structure and genetic variation in a non-redundant KP dataset. (a) Maximum-likelihood phylogenetic tree of the non-redundant KP dataset based on core SNPs, with a color gradient showing PC1 variation. (b) PCA of the non-redundant KP dataset based on LD-pruned SNPs (see Methods). Colors indicate geographic origin, while marker size and shape represent resistance and virulence scores, respectively, determined by Kleborate[23]. The sky-blue shading represents the density of PC1 values. The inset scree plot shows the variance explained (PVE) by the first 20 principal components.

The backbone of the KP tree is a line from which multiple branches radiate at different points along its length. This structure is suggestive of a continuously varying trait. Informally, the backbone has a top and a bottom, which can be seen in Figure 1a. The strains that branch at the top could, in principle, be given maximum backbone scores and those at the bottom minimum ones, with intermediate strains given values according to how far along the line they branch. We have not defined scores in this way, because there is a more systematic way of describing this variation, namely Principal Components Analysis (PCA)[21,22].

PCA identifies axes of variation that together explain the overall diversity within the species, with the first component, PC1, explaining the largest fraction of variation. When PCA was applied to the non-redundant global KP core genome SNP dataset used to construct the tree in Figure 1a, the first component (PC1) explained 16.7 % of the variation, with subsequent components each explaining approximately 8% or less (Figure 1b). PC1 captured the genetic variation underlying the phylogenetic backbone of the species, since strains with extreme PC1 values are found at either end of the backbone, while strains branching from the middle have intermediate values (Figure 1a). Strain loadings for PC1 are approximately continuous, albeit with a paucity of intermediate values (Figure 1b).

PCA is sensitive to all signals of relatedness, including clonal structure and can be prone to overfitting[24], especially where there are orders of magnitude more genetic markers than strains[24]. However, we found that PC1 was reproducible in different analyses of non-redundant datasets (Figure 2). PC1 strain loadings were consistent if components were defined by strains on the right-hand side of the phylogenetic tree (Supplementary Figure 3) or those on the left, implying that they are not generated by clonal structure (Figure 2a & b). PC1 values obtained when performing analysis on one half of the chromosome were correlated with those of PC1 calculated using the other half, implying that the signal is not due to specific genetic loci (Figure 2c & d). Specifically, they were also consistent when calculated using SNPs with extreme PC1 loadings or loadings that are closer to 0 (Figure 2e-h), implying that the signal affects genetic variation genome-wide.

**Figure 2.**
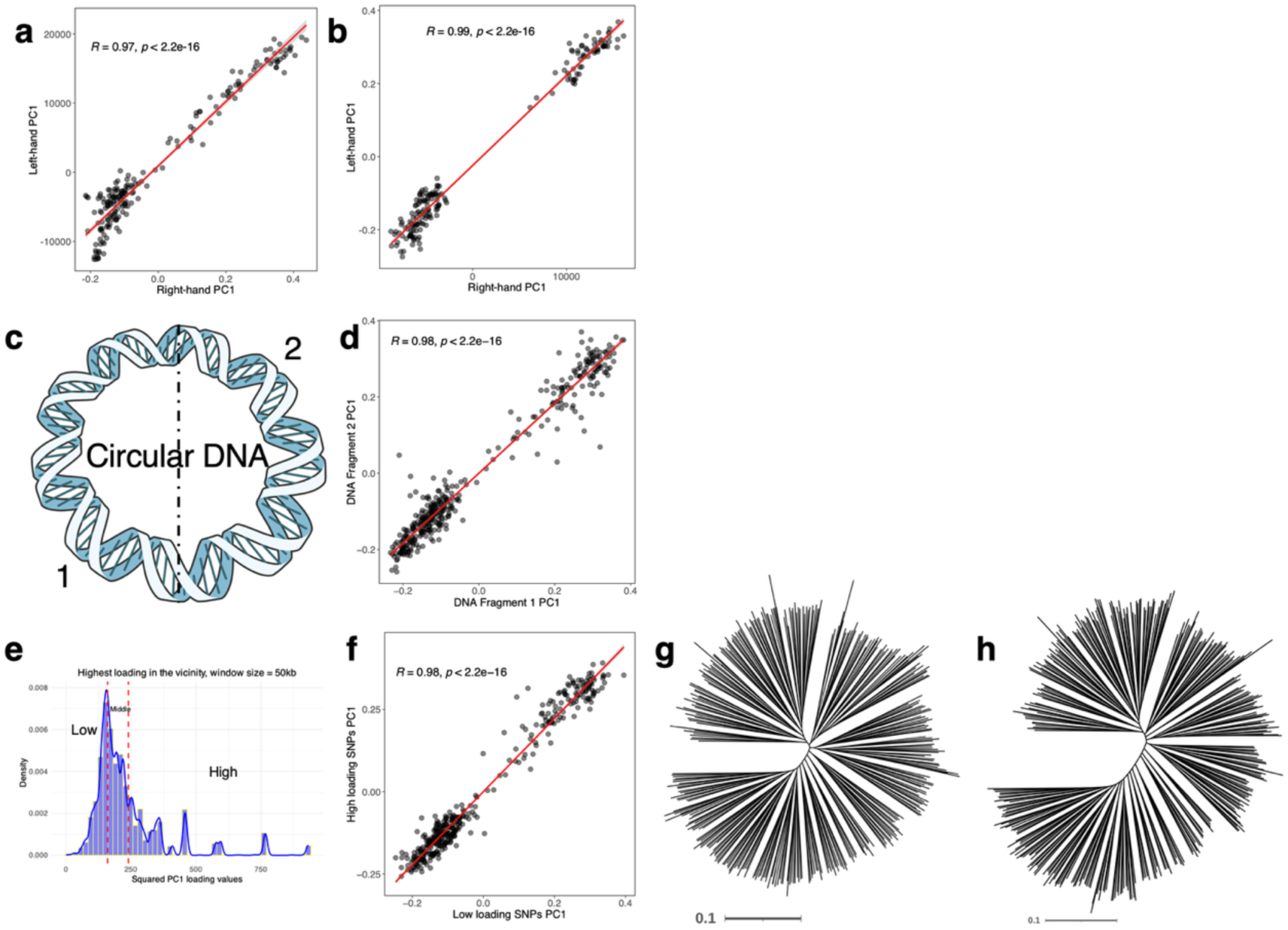
Three types of half-matching tests for the non-redundant KP dataset. (a–b) Half-strain matching: (a) Correlation of PC1 from strains on the right-hand of the tree projected onto left-hand group results. (b) Correlation of PC1 from the left-hand group mapped onto right-hand group results. (c–d) Half-genome Matching: (c) The genome is divided into two fragments: Fragment 1 (1 bp–2.6 Mb) and Fragment 2 (2.6 Mb–5.2 Mb). (d) Correlation of PC1 between the two fragments. (e–h) Partial-variants matching: (e) Histogram of SNP loadings on PC1, dividing SNPs into high- and low-loading groups. (f) Correlation of PC1 between high- and low-loading SNP groups. (g) Phylogenetic tree based on low-loading SNPs. (h) Phylogenetic tree based on high-loading SNPs.

To benchmark these results, we applied the same methods to data from a simulated neutrally evolving bacterial population (Supplementary Figure 4a & 4b), which will be discussed in more detail below. In the simulated data, PC1 did not explain as much of the variance as in the KP data (Supplementary Figure 4c). Furthermore, only one of three half matching tests gave significant correlation when applied to this data (Supplementary Figure 4c-i), contrasting with the results for KP. Thus, according to these criteria, we can falsify the simplest neutral models based on these results.

*Vibrio parahaemolyticus* (VP) also represents a useful benchmark for half-matching tests because its genetic structure is well characterized. The species has geographical population structure[19], which arises because the species lives in coastal waters and there have been barriers to movement of bacteria between oceans, albeit disrupted by recent human activity. VP also has a differentiated ecospecies “Molassodon”, which has nearly fixed differences with other “Typical” strains in approximately 40 core genes, while being undifferentiated in the rest of the genome[20,25]. This pattern is consistent with frequent recombination between ecospecies homogenizing variation in most of the genome, while natural selection has created and maintained differentiation at ecospecies loci.

When three different sorts of half-matching tests were applied to a global non-redundant dataset of, all gave reproducible results that reflected the population structure in the dataset (Figure 3a-f & Supplementary Figure 5a-e). However, when analysis was restricted to the Asian population (VppAsia), the phylogenetic tree is close to being a perfect circle and PC1 instead reflected ecospecies structure (Figure 3g-l & Supplementary Figure 5f-h). This structure generated weaker correlations in two of the three tests (Figure 3j and Supplementary Figure 5h), but there was no correlation between low loading and high loading loci, consistent with a single randomly mixing gene pool for most the genome[20,26]. Thus, the reproducibility of PC1 in all three tests in KP shows that the structure is genome-wide and in this respect is more like geographic population structure and less similar to ecospecies structure, which is only present in a fraction of the genome[26].

**Figure 3.**
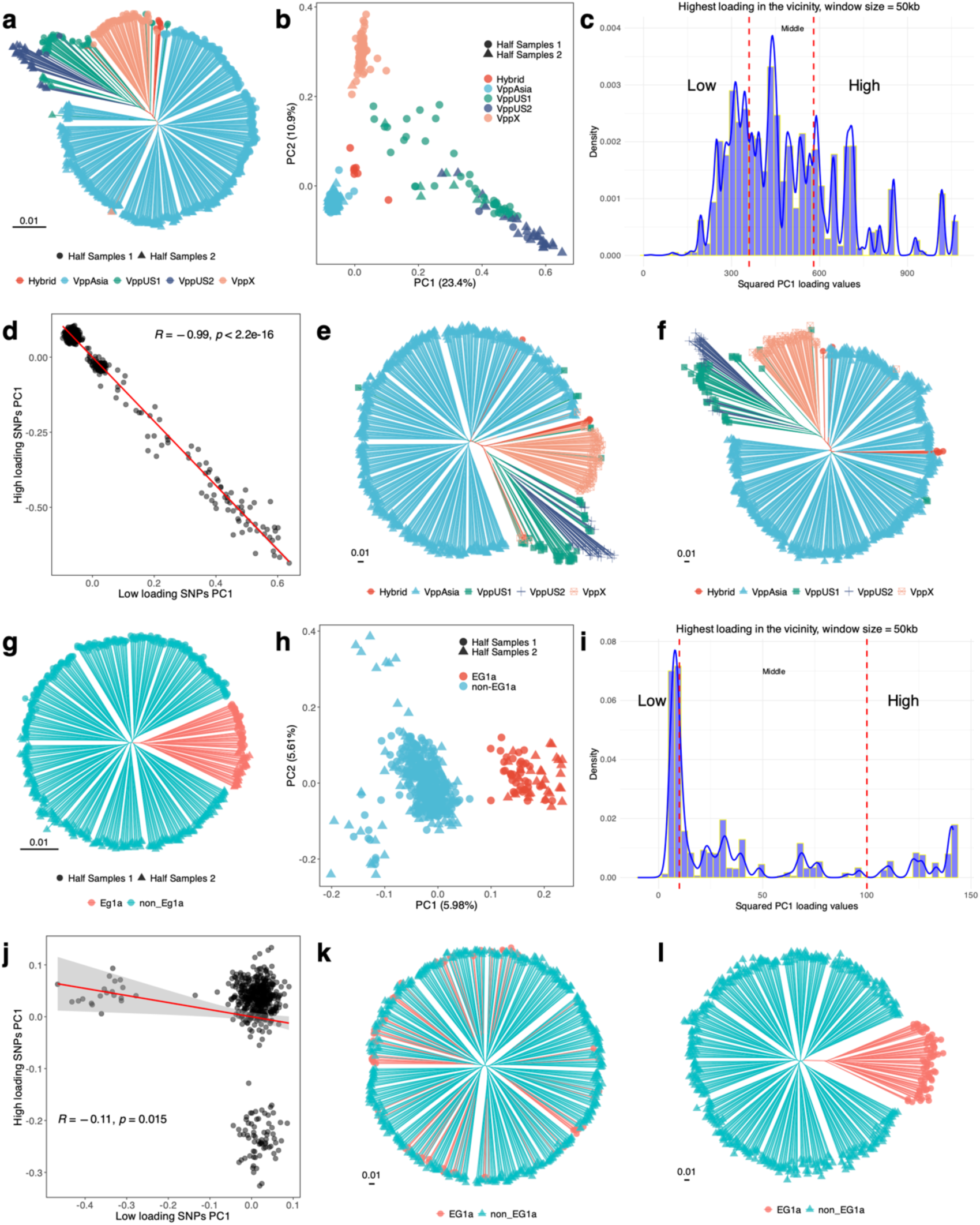
Partial-Variants Matching for *Vibrio parahaemolyticus* datasets. (a) Phylogenetic tree of the non-redundant VP dataset. Branch colors indicate populations defined by fineSTRUCTURE, and tip nodes represent the half-matching-based dataset partition. The Half-Strain Matching and Half-Chromosome Matching results are shown in Supplementary Figure 5 (b) PCA of the non-redundant VP dataset, using the same legend as in panel a. (c) Distribution of SNP loadings, with SNPs categorized into low, medium, and high groups based on the maximum loading value within 50 kb. (d) Correlation of PC1 between high- and low-loading SNP groups for the non-redundant VP dataset. (e) Phylogenetic tree constructed using low-loading SNPs for the non-redundant VP dataset. (f) Phylogenetic tree constructed using high-loading SNPs for the non-redundant VP dataset. (g) Phylogenetic tree of the VppAsia dataset. Branch colors indicate ecospecies, and tip nodes represent the half-matching-based dataset partition (see Supplementary Figure 4). (h) PCA of the VppAsia dataset, using the same legend as in panel g. (i) Distribution of SNP loadings, categorized into low, medium, and high based on loading values. (j) Correlation of PC1 between high- and low-loading SNP groups for the VppAsia dataset. (k) Phylogenetic tree constructed using low-loading SNPs. (l) Phylogenetic tree constructed using high-loading SNPs.

### Loci most strongly associated with PC1 are localized in the genome

In addition to strain loadings (Figure 1b), PCA generates SNP loadings, indicating which genomic variants are most strongly associated with the component. SNP loadings for PC1 showed four clear peaks (Figure 4a, left). Each peak contained many SNPs within less than 10 kb (Supplementary Figure 6). We also calculated loadings for accessory gene presence-absence, based on a Panaroo-generated[27] pangenome of the non-redundant strains, and found many had high loadings (Figure 4a, right).

**Figure 4.**
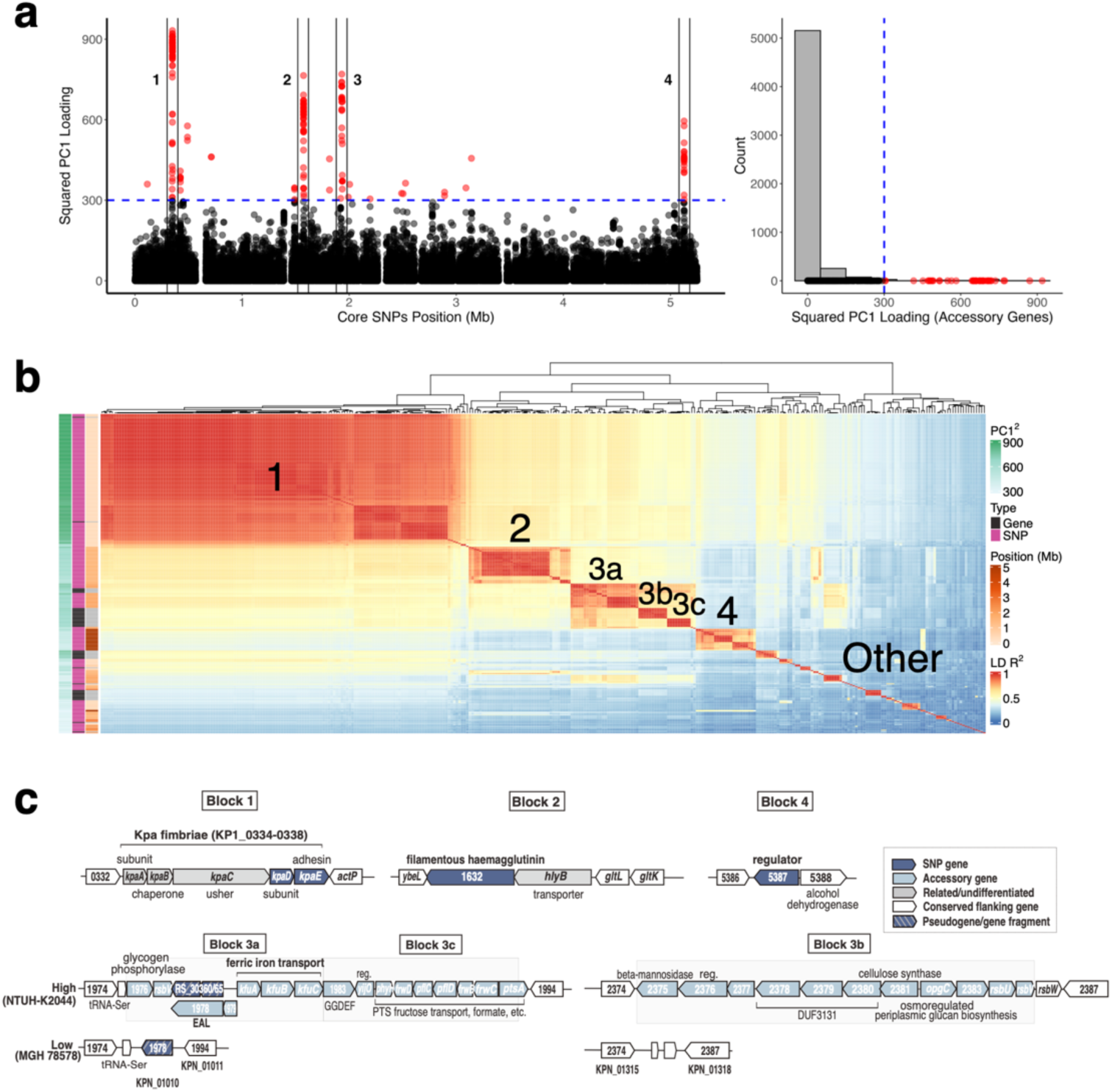
Location, Function, and Linkage Disequilibrium of PC1-Associated Loci. (a) Manhattan plot of squared core SNP (left) and accessory gene (right) loadings for PC1. Variants with values greater than 300 are defined as high loading variants and are marked in red. (b) Heatmap of pairwise LD *R^2^*values for high loading variants, with the first four blocks (including subblocks) labelled. (c) Functional annotation of blocks 1-4.

To investigate potential epistatic fitness interactions between the loci most strongly correlated with PC1, we chose a cutoff of 300 for squared PC1 loadings and calculated pairwise linkage disequilibrium (LD) values for all SNPs and genes above this cutoff. Hierarchical clustering of LD values organized the extreme loading variants into 44 blocks (Figure 4b). Notably, the top four blocks include a large fraction of all loci with PC1 values greater than 300, including the highest loading accessory genes. The organization and gene content of these blocks is shown in Figure 4c and Supplementary Figure 7.

Block 1 (Supplementary Figure 7a-c), which has the most extreme loading variants, includes two genes with SNPs (KP1_0337 & KP1_0338). These genes are part of the five-gene *kpa* fimbriae locus, which is part of the core genome in both KP and *K. variicola*[28,29], but found at lower frequency in other *Klebsiella* species. While the Kpa fimbriae are less characterized than those encoded by the *fim* locus, experiments in LM21 (a low PC1-loading strain) found that absence of *kpaC*, encoding the usher, significantly reduced *in vitro* biofilm formation[30]. The Δ*kpaC* mutant was not affected for binding to either a human cell line or *Arabidopsis* seedlings[30]. The most differentiated gene (KP1_0338/*kpaE*) encodes the adhesin subunit. The high and low variants are too distinct to have evolved within KP and cluster separately on a multispecies tree of the locus (Supplementary Figure 7c), implying that one or both variants was likely imported from another species.

Block 2 (Supplementary Figure 7d & e) and block 4 (Supplementary Figure 7f & g) also include SNPs in single genes (KP1_1632, filamentous haemagglutinin N-terminal domain-containing protein and KP1_5387, winged helix-turn-helix transcriptional regulator, respectively). The former, as for the *Kpa* fimbral adhesin, suggests a difference in adhesion, but so far appears uncharacterized. The regulator encoded by KP1_5387 in block 4 is also unstudied but seems always encoded next to an alcohol dehydrogenase in KP, *K. variicola*, *K. quasipneumoniae*, *K. africana*, *K. quasivariicola* strains.

Block 3 (Supplementary Figure 7h & i) includes both SNPs and accessory genes and can be subdivided into three sub-blocks (3a, 3b, 3c) based on LD patterns. The majority of strains fall into two configurations (Supplementary Figure 7g), with the high-loading configuration including all accessory gene elements and the low loading configuration including none, but there are also strains with intermediate configurations.

Block 3 genes are located in two different genomic regions in the reference strain, with block 3b genes encoded at a different location from block 3a and 3c genes (Figure 4c). Blocks 3a and 3c were previously highlighted as associated with invasive strains[31], as well as those implicated in bovine mastitis[32]. Key genes of block 3a are *kfuABC*, encoding a ferric iron ABC transporter, which aids iron acquisition and impacts mouse colonization[31,33]. The block also encodes two c-di-GMP-related proteins, a regulator, and glycogen/PTS (phosphotransferase) sugar utilization/uptake genes. Block 3c is adjacent to block 3a in the genome and contains genes related to sugar uptake, metabolism, and regulation (again via c-di-GMP). Block 3b contains accessory genes encoding a cellulose synthase, three DUF3131 proteins, OpgC (osmoregulated glucan production), and several regulatory proteins (hybrid sensor kinase, anti-sigma-factor and its antagonist).

### Functional enrichment analysis

To establish whether PC1 exerts a greater influence on core or accessory genome variation, we compared the distribution of absolute PC1 loading values between core SNPs, core genes and accessory genes. The dashed vertical lines indicate the mean loading value for each group, which are closely aligned, suggesting similar overall contributions to PC1 (Figure 5a). However, accessory genes display a longer right tail in the distribution, indicating the presence of more extreme loading values. This suggests that, although the average contribution is similar, a subset of accessory genes is more strongly influenced by PC1.

**Figure 5.**
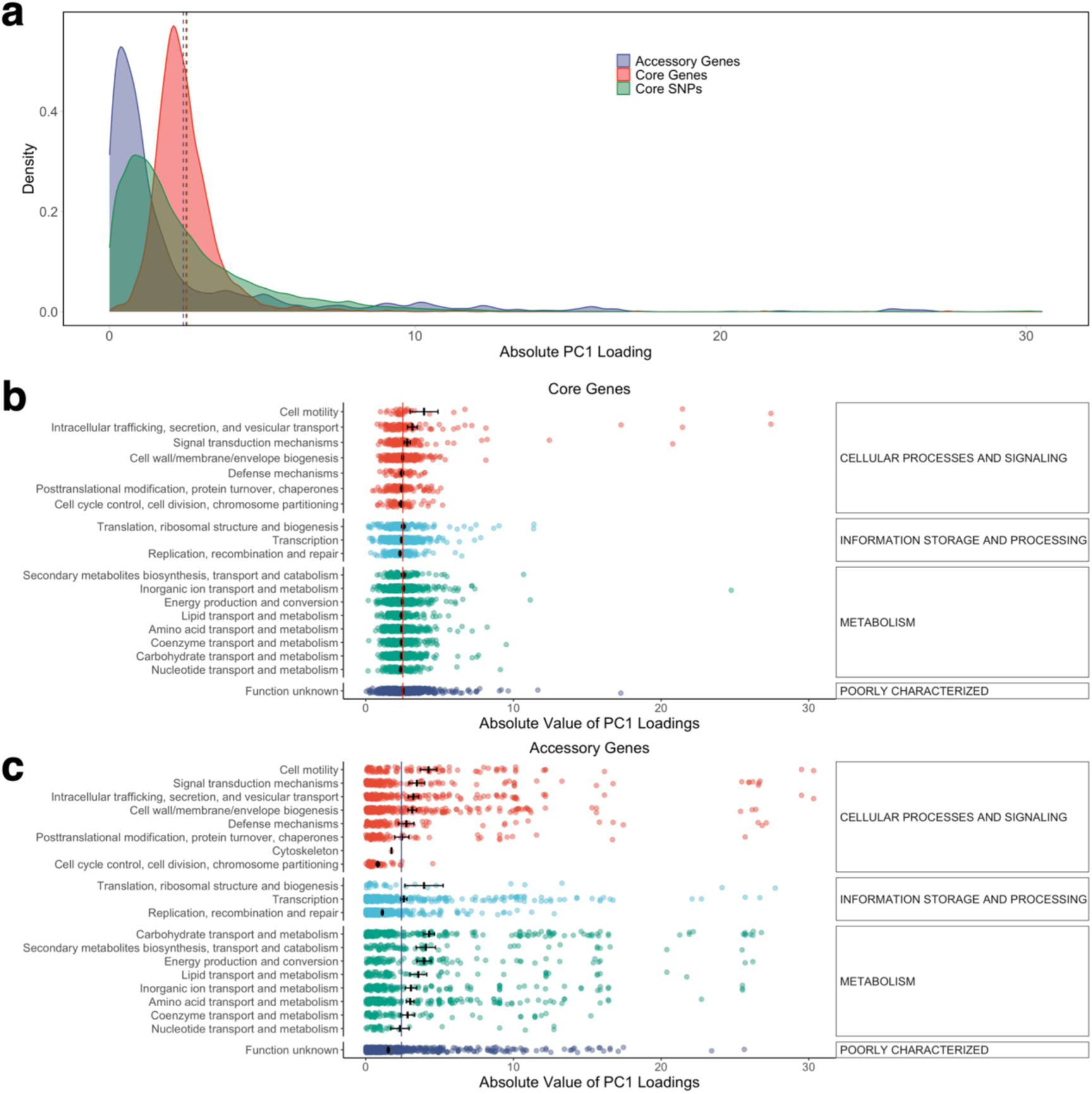
Overall and COG-specific distributions of absolute PC1 loading values for variants. (a) Density plot of PC1 loading values for core SNPs, core genes and accessory genes. Mean absolute PC1 loadings of core SNPs, (b) core genes and (c) accessory genes grouped by COG category. Bars represent the average absolute loading within each COG category, with error bars indicating the standard error. The solid (a) and dashed (b) lines mark the overall mean absolute loading across all features (genes or SNPs, respectively).

To establish whether there is a systematic relationship between gene function and SNP loadings, we calculated the mean absolute SNP loading for genes in each COG (Clusters of Orthologous Groups) functional category (Figure 5b) and found almost no difference between categories. However, larger and statistically significant differences were found for accessory genes (Figure 5c), with metabolic genes showing consistently higher mean absolute loading values than those involved in recombination and genome repair or uncharacterized genes.

### Virulence, antibiotic and isolation source profiles provide few clues

Although virulence and antibiotic resistance are important for the clinical assessment of KP, our analysis suggests that they are not the primary determinants of its overall population structure, which is not unexpected given that most lineages only cause human disease sporadically. Figure 1b shows the isolate metadata and Kleborate[23]-assigned resistance and virulence scores for the non-redundant dataset. Strains do not cluster based on geography, resistance, or virulence.

For multi-host bacteria, host-imposed selective pressures are important drivers of their evolution[34,35]. KP can be isolated from a variety of sources, including animals, plants, humans (which represent the largest proportion in databases), and water, including the sea. However, there is no strong evidence to support that host selective pressure is responsible for shaping the backbone of the KP phylogenetic tree (Supplementary Table 1).

Based on KP1_0338, the gene most strongly associated with PC1, we classified the multi-source strains into high and low variant groups (Supplementary Figure 8a). A Fisher’s exact test revealed a significant association between KP1_0338 variant group and host source (*p* < 0.01), primarily driven by strains from water and horses (Supplementary Table 1). However, we found that most water-isolated strains were from Manchester, where they were collected from patient or hospital wastewater sources. According to the Kleborate[23] annotation, most of these strains had a resistance score of 2 and a virulence score of 0 or 1, indicating that they are typical hospital-associated clones. We think this signal is an artifact of the strains mostly belonging to a handful of clonal lineages rather than evidence of true host specificity.

The KP isolates from horses were predominantly collected in France and Iceland, from various anatomical sites through both post-mortem examinations and live-animal sampling. However, when we projected the combined dataset — including the non-redundant dataset, along with strains isolated from water and horse sources — onto the results corresponding to Figure 1b (Supplementary Figure 8b), the projection revealed that the distribution of horse-derived strains was not significantly different from that of the non-redundant dataset. Thus, the results provide preliminary evidence for an association of the KP1_0338 locus with colonizing horses, but no evidence for a host association of PC1 as a whole.

In summary, our analysis finds no evidence that the observed population structure of KP is shaped by simple host-specific selective pressures. Moreover, no clear associations were observed between population structure and geographic origin, antimicrobial resistance, or virulence. Instead, the distribution of this gene appears to be largely independent of host origin, indicating that other ecological or evolutionary factors that are not captured by current metadata may underlie the observed patterns.

### The backbone is not explained by simple neutral models

The presence of this substantial component of variation raises the question: how has it arisen? Simple models of a bacterial population evolving with mutation and homologous recombination failed to recapitulate the patterns we observed in KP. For example, simulations with a population size of 2000 and one homologous recombination event per genome of size 10 kb approximately matched the overall LD decay curve seen in the real data (Figure 6a). However, for these parameters, there was no phylogenetic backbone evident, and PC1 failed to explain as much variance as in the real data (Figure 6b & c). As noted above, PC1 was also not reproducible when applied to different halves of the tree in a half matching test (Supplementary Figure 4b-e). More generally, we did not see a phylogenetic backbone in any simulations of a single neutrally evolving population (representative simulations shown in Supplementary Figure 9).

**Figure 6.**
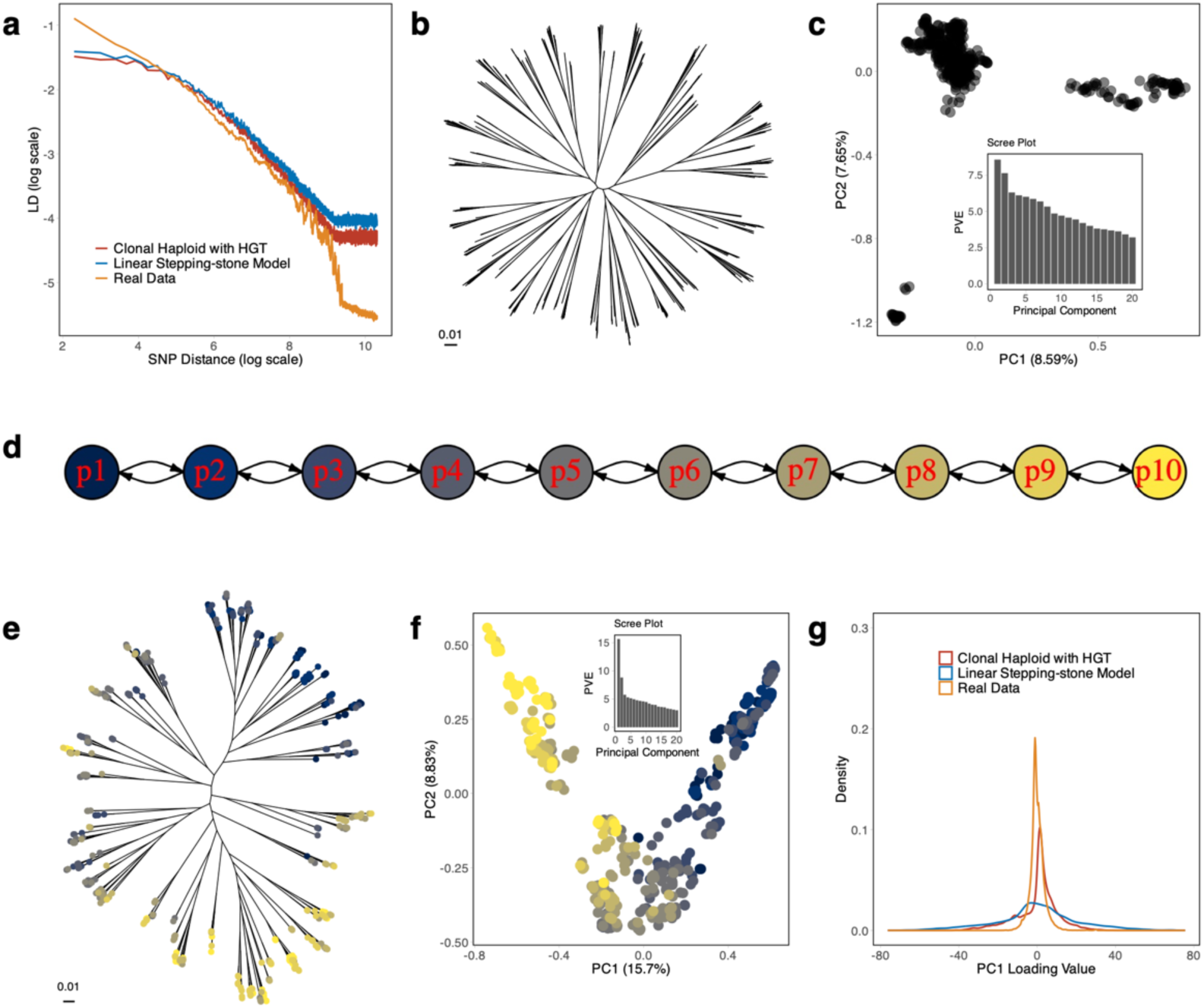
Simulated neutral models. (a) LD decay comparison to real data for best-fit neutral single population and ten-population stepping-stone models. Phylogenetic tree (b) and PCA results (c) for single population evolving neutrally. (d) Illustration of the ten-population stepping-stone model. Phylogenetic tree (e) and PCA (f) for the best fit stepping-stone model, coloured as shown in (d). (g) Density plot of SNP PC1 loading values illustrating the distribution of SNP loadings for real data and two neutral models.

The model of evolution used to generate the data in Figure 6a assumes that recombination rates are uniform across the species. More complex models are possible, for example in which recombination rates are high between closely related strains but lower between more distant ones. Such a pattern can arise for example if homologous recombination is suppressed by mismatches between donor and recipient sequence[36]. Such models are thought to explain much higher recombination rates within *Escherichia coli* phylogroups than between them[37]. Many models with these properties are possible, but since they all reduce the effectiveness of recombination, they are likely to make it difficult to recapitulate the low levels of LD in KP at large genetic distances. Moreover, the models that have been investigated previously cause species to cluster into discrete groups[38,39], rather than generating an approximately continuous pattern of variation. We are unaware of any model in which variable recombination rates within a single population predict a continuous structure similar to that seen in KP.

Genetic differentiation can arise due to geographical structure, or other factors that restrict gene flow between particular subsets of the population[20,26,40]. To explore the properties of geographical population structure we next simulated a stepping-stone model with 10 demes (Figure 6d). The demes are organised along a line, with a small number of migrants moving between adjacent demes each generation. Data simulated according to this model gave a tree with a phylogenetic backbone similar to that seen in KP, with PC1 explaining a similar fraction of the overall variation (Figure 6e, f and Supplementary Figure 11) and also approximately matched the LD decay curve (Figure 6a and Supplementary Figure 12).

However, the distribution of the SNP loadings for PC1 for these simulations did not match the real data, because there were many more SNPs with intermediate loading values (Figure 6g). Geographic population structure creates differentiation between populations through genetic drift, which is a random process that affects all of the genome but to differing degrees. The discrepancy in SNP loading values between simulated and real data implies that the variation in differentiation values in KP is higher than expected under a pure random drift model.

A longer-tailed distribution of drift levels, and hence PC1 values, could be generated by natural selection at a fraction of loci associated with adaptation to specific demes. However, even if a geographic model with some selection could be made to fit the data, this model seems to be poorly motivated in the case of KP. There is no evidence of KP being structured geographically (Figure 1b)[10,18] or of there being physical barriers to gene flow between ecological demes, since for example it is readily acquired by humans from a wide variety of ecological sources[41,42]. It is therefore hard to see how a model based on strong isolation between demes could be applicable to the species.

### KP variation is consistent with diversifying selection on a quantitative trait

Many traits are subject to diversifying selection[43]. For example, beak size has been shown to have two distinct optima in a species of Darwin’s finch, *Geospiza fortis,* which seems to have arisen because birds can specialize either on small soft seeds or larger nuts. Birds with intermediate beak sizes are not well suited to either type of food source and have lower fitness during droughts[44].

To investigate the signatures of diversifying selection, we performed simulations of selection on a single quantitative trait in a recombining bacterial population, in which strains with extreme trait values have higher fitness, while the fitness function is flat for intermediate trait values (see Methods). Simulation results in which half of all new mutations increased fitness by one and half decreased fitness by one showed that low recombination rates allow progressive divergence in trait values (Figure 7a-c). At intermediate recombination values, a phylogenetic backbone appeared that is similar to the one seen in the KP data (Figure 7 d-f), while high rates homogenize the population, leading to a circular star-shaped phylogeny (Figure 7g-i). For low and intermediate recombination rates (1 kb, 20 kb), PC1 values were strongly correlated with the true simulated trait values (Figure 7c, f). As expected, linkage disequilibrium values are strongly dependent on recombination rates (Figure 7k), with intermediate rates giving LD curves closest to the KP data.

**Figure 7.**
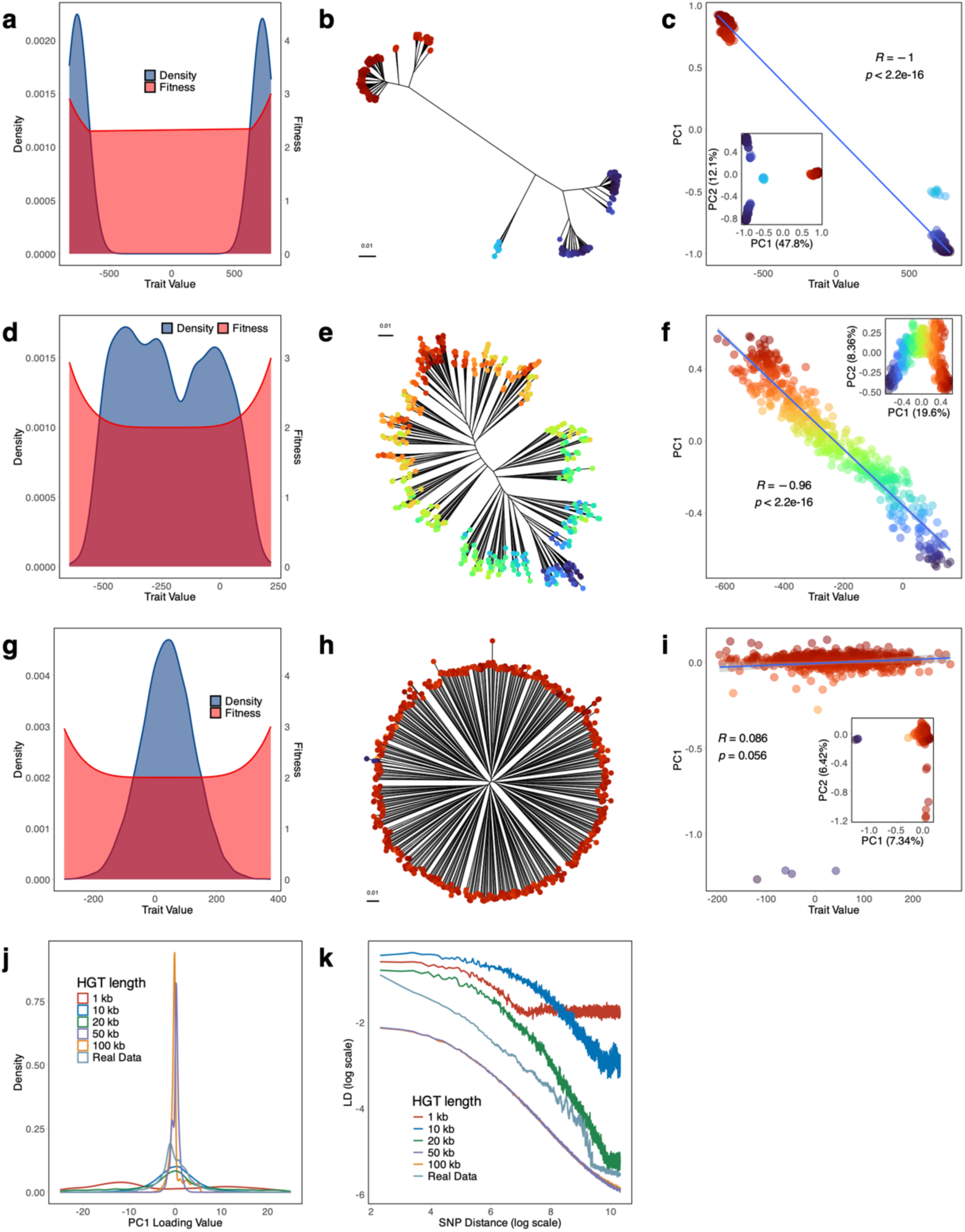
Simulation of diversifying selection on a polygenic trait at different recombination rates. (a, d, g) Trait value density distributions and fitness coefficients for recombination lengths of 1 kb, 20 kb, and 100 kb (Top to bottom) in final generation of simulation. (b, e, h) Phylogenetic trees depicting evolutionary relationships under each HGT length. (c, f, i) Correlations between PC1 values and trait values under each HGT length, with corresponding PC1 vs PC2 plots inserted in each panel. These simulations use 1Mb genomes. (j) Overall distribution of SNP loadings across all conditions. (k) Overall LD decay patterns across all conditions.

These simulations failed to recapitulate the sharp peaks in SNP loadings seen in the KP data (Figure 7j). We therefore modified the simulations by assuming that the distribution of the magnitudes of the effect of SNPs on the trait followed a Gamma distribution (Figure 8). High values of the Gamma shape parameter correspond to some mutations having a much larger effect on the trait than others. In these simulations, each PC1 loading peak corresponds to or is immediately adjacent to a SNP with a large effect on trait values. For high Gamma values, the number of peaks is smaller (Figure 8c, f & i).

**Figure 8.**
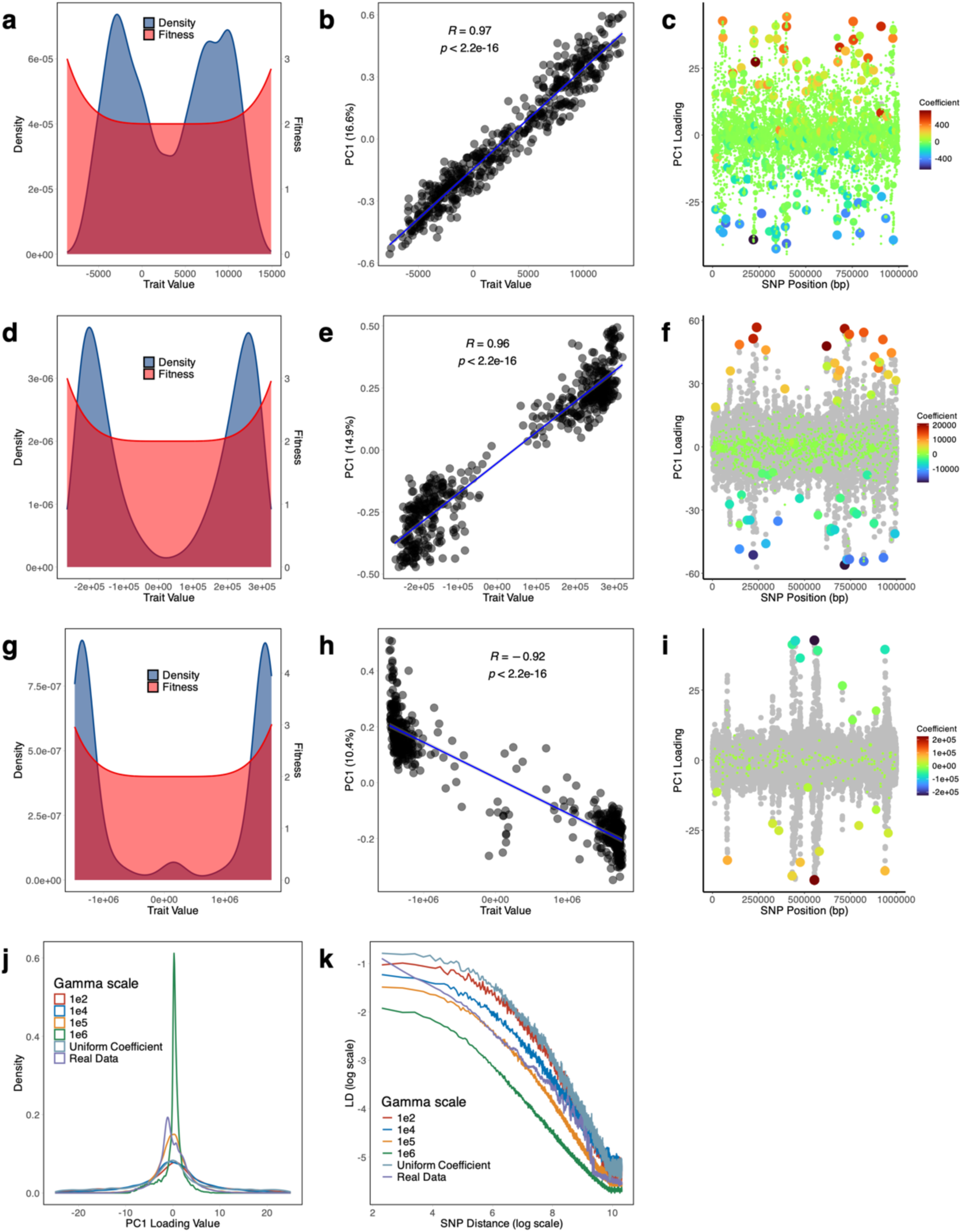
Effects of different selection coefficient on diversifying selection. (a) Distribution of fitness values under a gamma-distributed SNP effect size model (Gamma scale = 1e^2^), plotted against trait values. (b) Correlation between trait values and PC1 scores. (c) SNP loadings along PC1, with point colors indicating the magnitude of SNP coefficients. (d–f) Same as panels A–C, but under a gamma distribution with Gamma scale = 1e^4^. (g–i) Same as above, but with Gamma scale = 1e^5^. (j) Overall distribution of SNP loadings across all conditions. (k) Linkage disequilibrium (LD) decay pattern.

We found that simulations with a very large Gamma value could recapitulate the sharp peaks seen in the KP data (Figure 8j). For this parameter values, outside the sharp peaks, there is a great deal of variation in PC1 loadings, which is however not influenced by the SNP coefficients (Figure 8f & i) and therefore represents noise, that is generated by genetic drift. For high Gamma shape parameters, the correlation between strain PC1 values and the trait values accordingly becomes weaker (compare Figure 8h with Figure 8b&e), including if PC1 values are calculated only using top-loading SNPs (not shown). For intermediate Gamma shape parameters, the distribution of SNP loadings looked similar to that obtained assuming each SNP has the same effect on trait values (Figure 8j), but the distribution becomes more leptokurtic (with many SNPs with low values and long thin tails) as Gamma increases.

We found that a reasonable qualitative fit to the KP data could be found by simulating a 5 Mb genome length (similar to the true length) and assuming trait values for each SNP followed a Gamma distribution with a mean of 1 and a scale parameter of 5×10^6^ (shape of Gamma distribution = 2×10^-7^) (Figure 9). Specifically, we were able to recapitulate a phylogenetic backbone (Figure 9a & b), a PC1 correlated with the trait value (Figure 9e) and a small number of high loading peaks (Figure 9c), each of which is associated with SNPs with substantial influence on trait values. These simulations have slightly weaker LD than is seen in the real data (Figure 9h), with a smaller proportion of the variance explained by PC1. However, the pairwise LD between the highest loading SNPs showed a very similar pattern to that seen in real data (Figure 9d).

**Figure 9.**
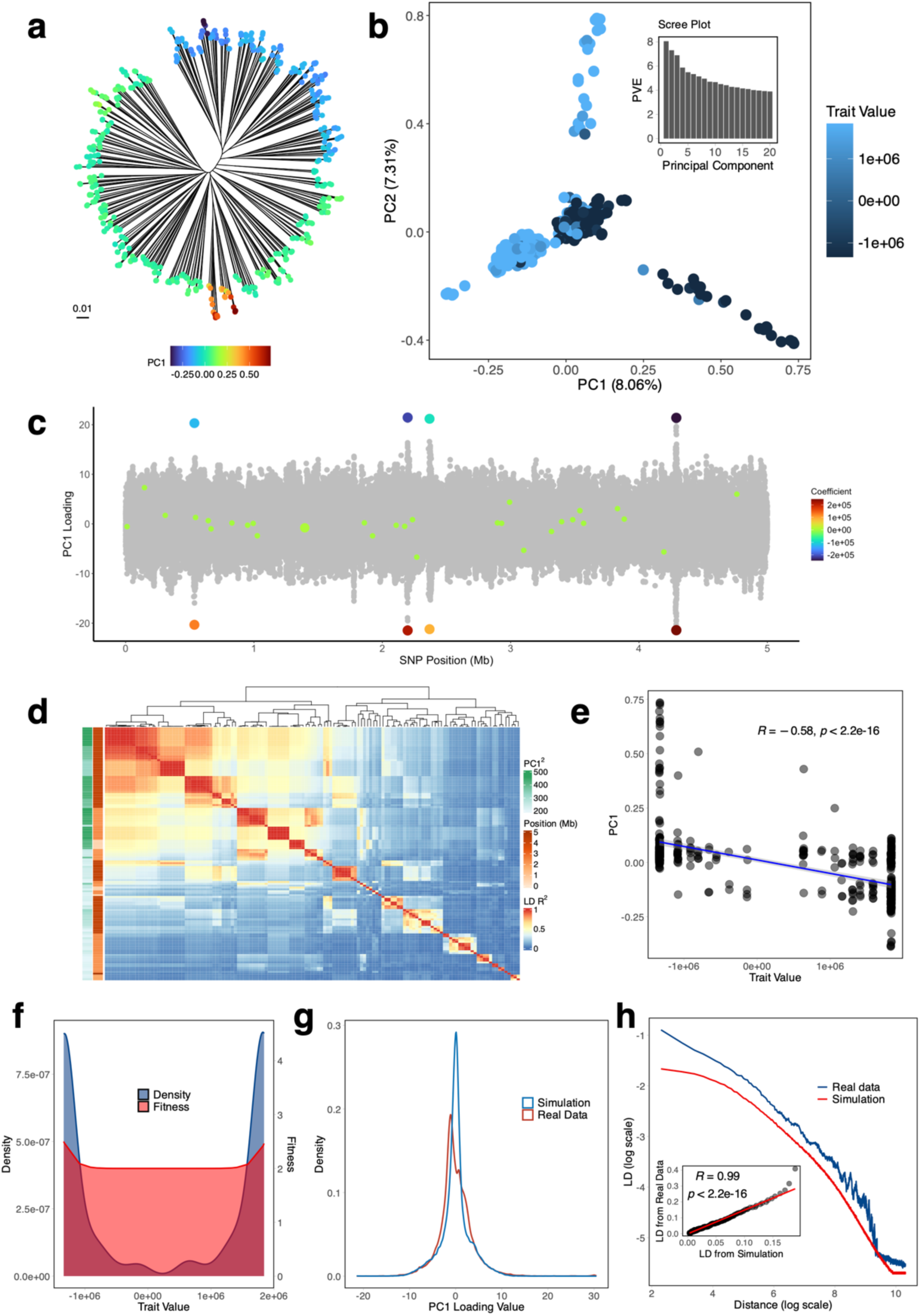
Best-fit simulated diversifying selection model. (a) Phylogenetic tree of simulated 5Mb genomes. (b) PCA results. (c) SNP loadings for PC1. (d) Heatmap of pairwise LD *R^2^* values for high loading SNPs. (e) Trait value density distributions and fitness function. (f) Correlation between trait values and strain PC1 values. (g) Distribution of PC1 SNP loadings for the 5 Mb simulation compared to the real data. (h) LD decay curves for the simulated and real datasets. An inset scatter plot compares the LD values between the two datasets, highlighting their correlation.

## Discussion

### Are quantitative trait models relevant for microbiology?

Quantitative trait models are central to our understanding of human disease genetics[45] and animal[46] and plant[47] biology. These models are applicable to many morphological and disease susceptibility traits, which fit the assumptions of the model, namely that the trait value is dependent on many variants, none of which explain a substantial fraction of the overall variation. Furthermore, each variants acts approximately independently of the others in its influence on the trait value[48,49]. These features allow the traits of offspring to be predicted based on the values found in their parents. The predictability inherent in these models has found several compelling practical applications, including predicting disease risk in humans using Polygenic Risk Scores[50] and to assess the genetic quality of livestock, based on estimated breeding values[51]. The same rules also make short-term evolution relatively predictable[52]. Quantitative genetics can be used for example to predict the change in beak sizes in populations of Darwin’s finches based on the intensity of selection and the additive genetic variation present within the population before selection takes place[44].

Bacteria do not undergo sexual reproduction in each generation, but instead reproduce clonally, while occasionally importing tracts of DNA from other strains[53]. The population-level response to selection depends on specific lineages rising in frequency[54]. As a result, the most straightforward applications of quantitative trait models in eukaryotes, such as predicting offspring trait values do not apply[55]. Despite these differences, bacteria do have many continuous-valued traits[56], such as cell size, growth rate and antibiotic minimum inhibitory concentration (MIC). It is of interest to understand the genetic variation of each trait circulating within species and how the trait responds to selection.

Laboratory selection experiments have provided important evidence on the genetic architecture of the response to selection in a novel environment[57]. Bacterial fitness in a particular environment is a continuous valued trait[58]. In asexual laboratory populations, mutations of large effect dominate the initial response, but diminishing returns epistasis sets in rapidly with later variants having progressively smaller effect[59]. Although these laboratory experiments provide excellent evidence on responses to novel environments for clonal populations, we currently have neither data nor theoretical models applicable to natural recombining populations of bacteria. For example, if a lineage is selected for increased cell size, we do not know whether the response is likely to be driven by variants of small or large effect, and whether the variants will principally arise *de novo* or be imported from other members of the population by homologous or non-homologous recombination.

To our knowledge, the only continuous valued trait whose variation has been studied extensively in bacterial populations is antibiotic MIC, which can be treated quantitively[60]. Antibiotic resistance is however an unusual trait, because of the episodic nature of exposure to antibiotics and the strong selection coefficients involved, which accentuates the role of variants of large effect[61]. The most important sources of variation are mobile genetic elements carrying antibiotic resistance islands and *de novo* point mutations[62,63]. These variants do not fit the assumptions of quantitative trait models because individual variants have a large effect and moreover, there are strong epistatic interactions between them[64,65]. In any case, when antibiotic selection pressure is applied, instead of increasing MIC across the population, the outcome is some lineages become highly resistant, while others remain susceptible[66].

One factor that is likely to be critical in determining the genetic architecture of a trait is the past selection history[67]. For example, if cell size has previously been consistently selected to be smaller within the natural environment, then we might expect the response to selection for larger cell size to be like that seen in laboratory selection environments[68,69], with *de novo* mutation dominating and the initial response generated by mutations of large effect. By contrast, if there has been fluctuating or diversifying selection in the population, then many variants affecting size may already be segregating. These existing variants will have already been tested by selection and will tend to be less disruptive to other features of the organism’s biology. Therefore, if we want to find traits that are polygenic within bacterial populations, where several variants behaving additively and most variants having small effect, we might focus on traits that have been under such selection regimes.

Here we have used simulation to demonstrate that there are circumstances in which selection on a quantitative trait might structure the genetic variation across an entire bacterial species. Specifically, we have performed simulations of an additive trait in which strains with the highest and lowest values of the trait have a small fitness advantage. In the simulations this tendency is counteracted by recombination, which randomises variation within the population. If the recombination rates at each locus are low, strains are likely to differentiate into two discrete groups with distinct trait values (Figure 7a-c). If recombination rates are higher, then intermediate strains can become common (Figure 7d-f) and variation becomes clinal. Because this cline is generated by selection on a trait, we call it a bacterial ecocline.

The clinal variation we see in our simulations depends on a balance between selection and recombination and can potentially be seen throughout the geographical range of the species. As such it is distinct from the clinal variation commonly seen in outbreeding animals and plants, which frequently has a geographical component. Eukaryotic biologists use the term ecocline to describe a situation where there is a single key ecological variable, such as altitude above sea levels for terrestrial animals or saltwater concentration for marine ones, which generates a strong geographical gene frequency cline in the loci that determine adaptation to that variable[70,71]. However, in their context, the gene frequency cline is dependent on limited dispersal between locations, which reduces the rate of recombination between individuals with high and low trait values. We think that our use of the term ecocline is more accurate, while theirs should strictly be called eco-geoclines. Their clines are, however, similar to the one we have simulated in being dependent on a key ecological trait.

### Clinal variation in KP

Fitness tradeoffs, such as between fast growth and starvation resistance are important for all bacterial species. However, given our limited understanding of selection in natural bacterial populations, it can be difficult to predict which traits might be subject to diversifying selection. We therefore suggest a “reverse ecology” approach [20,25,72,73] in which we first identify clinal genetic variation in bacterial species, before seeking to identify the trait or traits associated with the cline that are under diversifying selection.

Specifically, we noticed that the KP phylogenetic tree is structured by a backbone, which has been noticed informally by many researchers, but resists immediate explanation. To characterize the underlying variation in more detail, we used PCA. When PCA is applied to a non-redundant dataset, PC1 is strongly correlated with the backbone. However, caution must be exercised in interpreting PCA results, because the method will detect apparently clinal variation in essentially any dataset. The PC1 cline in KP is noteworthy because of the large fraction of variation it explains (Figure 1b), its reproducibility in analyses using subsets of the data (Figure 2) and the absence of geographic differentiation in the species.

Our PCA results are particularly useful because they give us the first clues about the genetic architecture of the clinal variation. Specifically, PC1 variant loadings have four prominent peaks, corresponding to LD blocks in the genome. In the simulations of selection on a quantitative trait that best fit these data, the Gamma parameter determining variation in trait values is large, meaning that most of the variation is generated by a small number of loci. In these simulations, we found that the highest PC1 loading peaks corresponded to the loci with the largest effect on trait values, but that remaining variation in PC1 appears to be approximately independent of trait values. That is to say loci with small effects on the trait had randomly distributed PC1 values, while most intermediate PC1 peaks correspond to neutral sites (Figure 8).

We have also visualized the linkage disequilibrium between these sites (Figure 1d) and found a pattern that is broadly consistent with the patterns seen in simulated ecoclines (Figure 7d-f). In our simulations of diversifying selection, aside from physical linkage blocks, most of the LD between sites is generated by selection on the trait, with pairwise LD values between unlinked blocks dependent on the amount by which each of the markers influences the trait. In the KP data, block 1 has the most extreme PC1 loadings, so that this block is predicted to have the largest effect on the putative quantitative trait. Concordant with this prediction, it has consistently high pairwise LD with loci in other blocks, while blocks 5-44, which have lower loading values, have consistently low pairwise LD values with other blocks. Very similar patterns are seen in the simulated data (Figure 7k). The LD patterns that we see thus seem broadly consistent with selection being mediated by the effect on a trait, rather than direct pairwise interactions between loci, which would generate a more patchwork-quilt like pattern.

### Phenotypic and ecological hypotheses about KP variation

Our hypothesis that selection is organizing KP variation into a bacterial ecocline entails two sub-hypotheses. The first is an ecological hypothesis — that diversifying selection on a trait has generated clinal variation — and a functional hypothesis — that the genetic variants showing clinal variation are modifying the selected trait. Demonstration of an ecocline requires validation of both hypotheses.

Existing functional annotations of the genes in the top four blocks provide hints, but not strong evidence that the genes might influence the same trait. The most differentiated gene is the adhesin component of Kpa fimbriae, which suggests the trait may be related to adhesion/diffusion. Block 2 encodes a filamentous hemagglutinin, which may also be an adhesin. The EAL- and GGDEF-domain proteins ecnoded in blocks 3a and 3c, respectively are related to c-di-GMP metabolism, a second messenger that controls trade-offs between sessile/surface and motile/free-living behaviours[74]. The accessory *kfuABC* genes in block 3a suggests there is some involvement of iron acquisition[33]. While these genes were previously found to be over-represented in invasive clinical isolates and bovine mastitis strains[32], we could not find any evidence that the trait is directly and specifically involved in human infection. One thing that is clear is that virulent or resistant genotypes can assemble on all positions of the ecocline.

We also attempted to identify specific gene function categories that were particularly associated with the backbone. For core genes, there was no difference between COG categories in mean absolute PC1 loading values (Figure 5). These results are consistent with our simulation results, which imply that if a trait is determined principally by a small number of loci, then genes that modify the trait by smaller amounts cannot be distinguished from neutral loci based on PC1 values. Therefore, it seems likely that outside the four peaks, most of the variation in PC1 values in core genome SNPs is generated by genetic drift. However, for accessory gene presence-absence, metabolic and cellular process-associated genes have systematically higher absolute loadings than genes of unknown function or those associated with replication, recombination or transcription, which provides suggestive evidence that these are under stronger selection.

Despite the provisional nature of the current evidence, we have been emboldened to propose KP variation might be organized into an ecocline based on our previous finding of "ecospecies" in two others bacterial genera[20,25,26] An ecospecies is like an ecocline, in that genetic differentiation is maintained by selection, in the absence of other barriers to genetic exchange. However, in contrast to bacterial ecoclines, each strain belongs to one ecospecies or another and there are no intermediates. For example, differentiation between Hardy and Ubiquitous *Helicobacter pylori* is largely restricted to 100 genes, which each have nearly fixed differences in at least 5 SNPs and there are no intermediate genotypes[26]. In VP differentiation has taken place at 40 core genes, and once again there are no intermediate genotypes[20].

For ecospecies, we have also been able to make stronger functional predictions, relating the VP ecospecies to motility[20] and the *H. pylori* ecospecies to host diet[26]. We have also recently used laboratory experiments to confirm and extend the original functional predictions and show that the differentiation between VP Typical and Molassodon ecospecies is functionally coherent, with the Molassodon having evolved a hunt, kill and devour phenotype that it employs in viscous liquids[25]. These results show that the reverse ecology approach can identify loci that act together in natural populations.

The absence of equivalent functional predictions means that our current designation of a bacterial ecocline in KP is provisional and needs to be validated by phenotypic data. If the ecological hypothesis is correct, and the genes with extreme loadings are influencing a single trait, then characterization of the effects of these variants should simultaneously provide considerable insight into gene function, ecology and evolution for a key bacterial fitness trait. This would likely represent a breakthrough as these aspects are hard to study individually, handsomely justifying the initial effort required to identify the trait.

It should be cautioned however that diversifying selection may not reflect a single measurable trait. For example, distinct foraging strategies in Darwin’s finches might require different beak sizes but also different behaviors, which may also be genetically determined. This could create linkage disequilibrium between genes determining beak size and foraging behavior. This ecologically-generated linkage disequilibrium is also evident in the Molassodon ecospecies example[25], where genes responsible for motility, bacterial killing and nutrient uptake have coevolved. In these cases, it might be hard to find a laboratory condition that links together the function of the different genes involved in the ecocline.

A challenge and virtue of the reverse ecology approach[25,72,73] is that it confronts us with the gulf between laboratory conditions and the natural environment that we seek to understand, and more specifically our difficulty in putting ourselves in the same shoes as bacteria and predicting the ecological challenges they are likely to face. In any case, we propose that investigation of phenotypic differences associated with the PC1 cline has the potential to deliver key insights into KP ecology, as well as on the biology of quantitative traits in microbes.

## Methods

### Genome collection and sequence alignment

We collected 1,421 KP genomes from the NCBI database, selected according to iterative-PopPUNK test dataset^78^, along with a reference genome (KP strain NTUH-K2044, NC_012731.1). Additionally, we obtained genome metadata using the NCBI Datasets command-line tools (v16.12.1)[76] and used Kleborate (v2.4.0)[23] to extract clinically relevant genotypic information on strains. All genomes were aligned to the NTUH-K2044 (NC_012731.1) reference genome using Nucmer (v3.1)^61^ for reference-based alignment.

In total, 8,331 global VP genomes were downloaded from the NCBI database, based on the dataset from the study ’Why panmictic bacteria are rare’[77] and aligned to the VP RIMD 2210633 (RefSeq NC_004603.1 and NC_004605.1)[78] using Nucmer (v3.1)[79], following the approach described in previous VP studies[19,20].

### Core genomic variant calling and definition of non-redundant dataset

SNP-sites (v2.5.2)[80] was used to call all the variants from the whole KP dataset alignment. Core SNPs were defined as the SNPs present in at least 99% of the genomes across the whole genome alignment, resulting in 850,158 core SNPs after extraction. Using these core SNPs, pairwise distances were calculated across all genomes with snp-dists (v0.8.2)[81]. A non-redundant dataset of 369 genomes was defined by selecting genomes with pairwise SNP distances >20 kb (Supplementary Table 2).

The same method to the whole VP dataset, using a 50 kb SNP distance threshold to reduce redundancy, yielding 667 genomes (Supplementary Table 3) with 1,619,597 core SNPs.

### fineSTRUCTURE and population assignment

To further investigate the population structure of the non-redundant 369 KP strains described above, we performed chromosome painting and fineSTRUCTURE analysis. Using ChromoPainter mode in fineSTRUCTURE (v4.1.0)[82], we inferred chunks of DNA donated from a donor to a recipient for each recipient haplotype and summarized the results into a co-ancestry matrix. Subsequently, fineSTRUCTURE clustered 369 individuals based on the co-ancestry matrix, with 100,000 iterations for both burn-in and Markov Chain Monte Carlo (MCMC). In this analysis, only variants present in all non-redundant strains were considered, as fineSTRUCTURE requires input data without missing values.

To assign a population to the non-redundant VP dataset, we followed the same fineSTRUCTURE approach using haplotype data from core SNPs. Based on the fineSTRUCTURE results, the genomes were classified into five populations: VppAsia (506 strains), Hybrid (7 strains), VppUS1 (50 strains), VppUS2 (30 strains), and VppX (74 strains). Within the VppAsia dataset, strains were further categorized into ecospecies types (EG1a and non-EG1a) based on 27 differentiating genes[20].

### Construction of phylogenetic trees

Based on the core SNPs defined above, we constructed nucleotide sequence alignment files for both the entire KP dataset and the non-redundant KP dataset. Phylogenetic trees were constructed using FastTree (v2.1.10)[83] with the "-nt" option under the default model. The trees were visualized and compared using the ggtree package (v3.12.0)[84] in R.

Similarly, we used the same software and parameters to construct phylogenetic trees for the non-redundant VP and VppAsia datasets, as well as the simulated datasets. During Partial-Variants Matching, we also constructed phylogenetic trees based on a subset of SNPs from the real datasets.

### Principal Component Analysis

Before performing PCA (Principal Component Analysis) on the non-redundant KP dataset, PLINK (v1.9)[85] was used to filter out loci with a minimum allele frequency (MAF) below 2% from the core SNPs, leaving 102,226 variants. Linkage disequilibrium (LD) pruning was subsequently performed to remove linked SNPs, where one SNP from each pair was removed if the LD (*R^2^*) exceeded 0.1 within a 50 kb window. The window was then shifted forward by 10 bp, and the procedure repeated to ensure comprehensive coverage. A total of 33,589 pruned variants were obtained and used for conducting PCA on the non-redundant dataset with PLINK[85], extracting and reporting the first 20 principal components by default.

Similarly, we used the same software and parameters to perform PCA on the non-redundant VP dataset, which included 220,737 SNPs after MAF filtering and 69,537 SNPs after pruning. We also applied PCA to the VppAsia dataset, containing 207,292 SNPs after MAF filtering and 65,058 SNPs after pruning. For the simulated datasets, we only applied MAF filtering to retain enough SNPs, followed directly by PCA without pruning.

### Variant loadings calculation

We first calculated the correlation between the original variables and the principal components, which is specifically referred to as the PCA loading for strains. The loadings can be determined as the coordinates of the variables (eigenvectors) multiplied by the square root of the eigenvalues associated with that principal component (Equation 1). When performing PCA, PLINK[85] generates output files ending with eigenval and eigenvec. Next, we transform the MAF-filtered SNP matrix into a binary matrix, assigning a value of 1 to minor alleles and 0 to all others (including the third and fourth most frequent alleles at non-biallelic sites). Finally, through matrix multiplication, we computed the product of the core SNP matrix and the PCA loading matrix to obtain the loading values for the MAF-filtered SNPs (Equation 2).

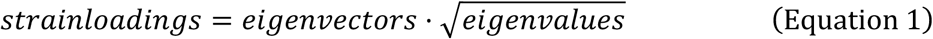

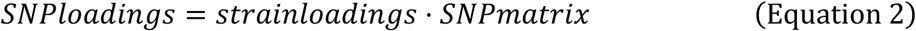

To better highlight the significant peaks in the non-redundant KP result, we plotted the squared PC1 SNP loading values and marked the dots with squared loading values greater than 300.

Similarly, we used the same method to calculate SNP loadings for the non-redundant VP dataset, the VppAsia dataset, and the simulated datasets.

### Pangenome analysis

To estimate the gene content, we annotated all KP genomes with prokka (v1.13)[86] software, using a curated version of the NC_012731.1 (NTUH_K2044) proteome as the primary annotation source. The Generic Feature Format (GFF) outputs generated by prokka were then used as the input files for panaroo (v1.2.8)[27] with the strict model. The "--remove-invalid-genes" and "--merge_paralogs" options were also employed during the analysis. Next, we extracted the non-redundant dataset from the overall results and defined 5,558 accessory genes as those present in between 2% and 98% of the genomes in the non-redundant dataset. Using the same approach (Equation 2), we also calculated the gene loadings and defined genes with squared PC1 loadings greater than 300 as high-loading genes.

### Calculation of linkage disequilibrium decay

To analyze the decay of linkage disequilibrium (LD) in KP, we included 102,226 MAF-filtered SNPs and used the Haploview (v4.2)[87] software to compute *R^2^* values. When running Haploview, we set "-maxdistance 30" to ensure that the maximum marker distance for *R^2^* calculation did not exceed 30 kb. Additionally, we set "-minMAF 0" and "-hwcutoff 0" to exclude any filtering based on minor allele frequency and Hardy-Weinberg equilibrium testing, ensuring that all input SNPs were included in the calculations. The average *R²* value was calculated using a 10 bp step size.

Similarly, the same software and parameters were used for the simulated datasets to obtain the LD decay results for the simulated data.

We also calculated the LD between the high-loading variants defined above (340 variants in total, including core SNPs and accessory genes) used Equation 3 in R. Each element 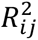 of the matrix quantifies the degree of linkage disequilibrium between variant *i* and variant *j*, providing a comprehensive view of the genomic associations among core SNPs and accessory genes. Based on manual inspection, we removed an accessory gene (group_205) that represented fragmented versions of KP1_1982 (339 variants left), which is visualized in a heatmap, generated using the ComplexHeatmap package[88].

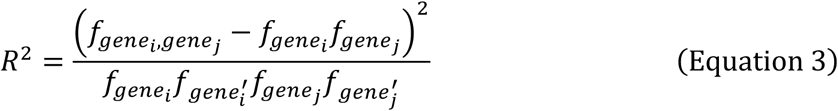

Based on the LD heatmap of all high-loading variants, we performed hierarchical clustering to divide the heatmap into 44 blocks. Among these, four larger blocks contain multiple variants, while the remaining 40 smaller blocks each contain only a few variants. Notably, block 3 was further subdivided into 3a, 3b, and 3c based on the physical locations of the mutations on the reference genome. We observed that some SNPs in block 3 are located within the KP1_RS30360 and KP1_RS30365 genes in the reference genome. According to the annotation file of the reference genome, KP1_RS30365 is classified as a pseudogene. However, we do not concur with this interpretation. We believe that KP1_RS30360 and KP1_RS30365 together represent a complete gene, with some strains (*e.g.*, MGH 78578) containing only a fragment of this gene. the differentiation at this locus also likely represents a functional presence/absence rather than functional differentiation via SNPs.

As mentioned above, we converted the variants into binary, where 0 indicates that the core SNP is a major SNP and the accessory gene is absent in the strain, and 1 indicates that the core SNP is a minor SNP, and the accessory gene is present in the strain. We then generated heatmaps of the variant states according to the blocks, with rows following the same order as the LD heatmap and columns subjected to hierarchical clustering.

### Phylogenetic reconstruction of a gene across multiple *Klebsiella* subspecies

Genomes were retrieved from the Pasteur Institute Klebsiella MLST Database by applying the following filtering criteria: (i) the taxonomic designation must be one of the following subspecies: *K. quasipneumoniae subsp. quasipneumoniae*, *K. quasipneumoniae subsp. similipneumoniae*, *K. variicola subsp. variicola*, *K. variicola subsp. tropica*, *K. quasivariicola*, or *K. africana*; (ii) the annotation status for MLST, ribosomal MLST, and scgMLST629_S must all be labeled as good; and (iii) assembly quality checks including the number of contigs, assembly size, minimum N50, and %GC must all be pass.

To confirm species identity, the resulting genomes were further screened using Kleborate[23]. The gene KP1_0338 from the reference strain was used as a query in a BLAST search, and MGH 78578 was included as an additional representative strain. To avoid overrepresentation of any subspecies, 10 genomes were randomly selected from each subspecies identified in the BLAST results.

These selected genomes, together with the reference strain, MGH 78578, and the 5 strains with the highest and 5 strains with lowest PC1 loadings from the previously defined non-redundant dataset, were aligned using MAFFT[89]. The resulting multiple sequence alignment was then used to construct a phylogenetic tree using FastTree[83].

### Functional annotation

To further characterize the functional content in KP, we annotated both the MAF-filtered SNPs (MAF > 0.02) with SnpEff (v4.3t)[90], using the reference genome NTUH-K2044 (NC_012731.1), and applied custom filtering to focus on coding region variants by disabling annotation for upstream, downstream, and UTR (untranslated) regions. We then obtained the types of the SNPs and determined whether they were synonymous mutations. For accessory genes, we used the eggNOG-mapper[91] online tool with its default settings to ensure a comprehensive and standardized annotation.

For the identified high-loading variants (squared PC1 SNP loading > 300), we further mapped these variants to the reference genomes of KP strains MGH 78578 (NC_009648.1) using BLASTn[92]. The mapping aimed to identify their genomic locations and potential functional implications in these strains. Additionally, we searched the UniProt[93] database to retrieve gene names and associated functional annotations for the high-loading variants.

We calculated the mean absolute PC1 loading for accessory genes within each COG category, as well as the overall mean absolute loading across all accessory genes. For core SNPs, we first computed the mean absolute PC1 loading of SNPs belonging to each core gene, then calculated the average of these values for each COG category. In addition, we determined the overall mean absolute loading across all core genes.

We visualized the results using bar plots, where each bar represents the mean absolute PC1 loading of genes (core or accessory) within a given COG category, and the error bars indicate the standard error of the mean. A solid line denotes the overall mean absolute PC1 loading across all accessory genes (or core genes), serving as a reference.

### Isolation background characteristics

To investigate potential host specificity in KP, and to address the overrepresentation of human-derived isolates in our primary dataset, we incorporated a multi-host dataset from the Pasteur Institute *Klebsiella* MLST Database (https://bigsdb.pasteur.fr/cgi-bin/bigsdb/bigsdb.pl?db=pubmlst_klebsiella_isolates). Isolates were selected based on the following quality criteria: chromosomal assembly size between 5 and 6 Mb, fewer than 100 contigs, an N50 greater than 20,000 bp, fewer than 1,000 ambiguous bases (Ns), and a GC content between 55% and 60%. Additionally, only genomes with “good” annotation status for MLST, ribosomal MLST, and SCGMLST629_s schemes were included. Kleborate[23] was applied to refine species classification, and *Klebsiella pneumoniae* species complex (KpSC) isolates were excluded. To minimize overrepresentation of human-associated strains, genomes with host information explicitly annotated as “human” were excluded. However, a small number of human-derived isolates may still be present due to inconsistent or alternative host labels (e.g., *Homo sapiens*). (Supplementary Table 4).

We used the KP1_0338 gene from the reference strain as the query sequence and included MGH 78578 as an additional representative. These two strains represent the extremes along the PC1 axis in our population structure analysis, with the reference strain showing an extremely high PC1 value and MGH 78578 a low one, thus capturing the two major variants of KP1_0338 present in the KP population. Using BLASTn[92], we identified KP1_0338 homologs in the multi-host dataset. The retrieved sequences were aligned using MAFFT[89], and a phylogenetic tree was constructed with FastTree[83] to examine sequence diversity. Based on the topology of the KP1_0338 phylogeny, isolates were grouped into two distinct clusters, designated as High and Low types.

To assess host-specific enrichment of these gene variants, we performed a one-vs-rest Fisher’s exact test for each host. This test compared the counts of High and Low types within each host to those in all other hosts combined.

We projected the combined dataset—including the non-redundant dataset as well as strains isolated from water and horse sources—onto the principal component space defined by the non-redundant dataset. For each group (non-redundant, water-derived, and horse-derived), we calculated the mean PC1 loading values and visualized them with standard deviations as error bars. Pairwise comparisons between groups were performed using two-sample t-tests to assess the statistical significance of differences in PC1 loadings.

### Half-matching testing

To assess the stability and repeatability of PC1 in KP, we used three half-matching tests, each designed to evaluate the consistency of PC1 under varying conditions.

#### 1. Half-Strain Matching

Firstly, we divided the non-redundant KP dataset into two groups based on the backbone in the phylogenetic tree. We then performed PCA on the first group using the pruned SNPs. Afterward, we calculated the MAF-filtered SNP loadings using the above steps. Subsequently, we obtained the product for the second group through matrix multiplication, representing the mapping of the PCA results from the first group onto the second group. We then calculated the correlation for the PC1 between the mapped results of the first group and the PCA results of the second group. Conversely, the same matching analysis was performed on the data from the second group.

#### 2. Half-Genome Matching

KP has a circular chromosome with a length of approximately 5 to 6 Mb, and the reference genome we used is approximately 5.2 Mb in length. We divided the circular chromosome into two DNA fragments (Fragment 1 (1 bp–2.6 Mb) and Fragment 2 (2.6 Mb–5.2 Mb), each containing approximately half of the chromosome. Specifically, the first dataset comprised the pruned SNPs in the first half of the chromosome, while the second dataset encompassed the unlinked SNPs from the second half. We separately performed PCA on these two datasets using PLINK[85]. Finally, we calculated the correlation between the PC1 of the two fragments.

#### 3. Partial-Variants Matching

First, we assigned maximum loading values to the MAF-filtered SNPs based on their squared PC1 loadings, using a 50 kb window size with a 1 bp step size. This process identified the maximum squared PC1 loadings within each 50 kb vicinity. Based on the distribution of maximum squared PC1 loadings, the SNPs were categorized into three types: low-loading SNPs, medium-loading SNPs, and high-loading SNPs. We separately conducted PCA on the high-loading SNP dataset and low-loading SNP dataset, by extracting unlinked SNPs from each. Subsequently, we calculated the correlation between the PC1 of the 2 datasets and constructed the phylogenetic tree for the 2 datasets. In this test, we ignored SNPs with intermediate loading values.

Similarly, the three types of half-matching tests were applied to the non-redundant clonal population simulation (Clonal Haploid Bacteria Model with Horizontal Gene Transfer, SNP distance > 2 kb, shown in Supplementary Figure 4a), the non-redundant VP dataset and the VppAsia dataset. However, in the half-strain matching test, although we still performed splits along the phylogenetic tree, these datasets were not always divided into left-hand and right-hand subsets in the trees. Therefore, in these datasets, we referred to the two subsets as half samples 1 and half samples 2. Additionally, in VP, which contains two chromosomes, the half-genome matching is implemented by separating the data according to chromosome number rather than splitting each chromosome into two segments.

### Simulation of bacterial populations

We used SLiM (v4.0.1)[94] to simulate the evolutionary dynamics of bacterial populations under different scenarios using a non-Wright–Fisher (nonWF) framework[95]. All scenarios started with identical initial conditions: each bacterial genome consisted of a single 1 Mb haploid chromosome, with a mutation rate of 10^-6^ per site per generation, approximately ten times higher than the estimated mutation rate for KP^92^. Horizontal gene transfer (HGT) occurred at a rate of 1.0, ensuring that every individual experienced HGT in each generation. The simulations ran for 10,000 generations. Two mutation types, "m1" and "m2", occurred at each site with equal probability (1:1) and were assumed to be neutral. However, in the final simulation, we adjusted the chromosome length to closely match the length of a real KP genome.

### Modeling clonal haploid bacteria with horizontal gene transfer

In the first simulation scenario, we modeled the evolutionary dynamics of a single bacterial population, focusing on the spread of a single type of neutral mutation and the genetic exchange within population via HGT. The HGT tract length was set to 100 bp, 1 kb, 10 kb, or 100 kb, representing a range of recombination sizes to accurately capture genetic variation within the population. Each bacterium produced offspring according to a Poisson distribution with λ = 2, meaning that, on average, each bacterium produced two offspring per generation. However, the number of offspring per bacterium in any given generation could vary, with some bacteria producing more and others producing fewer, reflecting the probabilistic nature of the Poisson distribution. We conducted three trials with population sizes of 1,000, 2,000, and 10,000.

### Linear stepping-stone model

In the second simulation scenario, we constructed a linear stepping-stone model with 10 subpopulations of clonal haploid bacteria, where each subpopulation was connected by migration only to its nearest neighbors in the chain. The average recombination tract length within each subpopulation was set to 10 kb. Migration occurred with varying numbers of migrants — 1, 10, 20, or 50 individuals — moving to neighboring subpopulations each generation to simulate gene flow driven by geographical factor or other isolating barriers. Each individual produced offspring according to a Poisson distribution with λ = 2. We conducted three trials with total population sizes of 1,000, 2,000, and 10,000, corresponding to 100, 200, and 1,000 individuals per subpopulation, respectively.

### Selective model

In the third simulation scenario, we focused on a population of 10,000 clonal haploid bacteria, where generations 1–5000 evolved neutrally, and selective pressure was introduced from generations 5001–10,000. Horizontal gene transfer (HGT) tract lengths were set to 1 kb, 10 kb, 20 kb, 50 kb, and 100 kb. In this model, the coefficient for any locus on the chromosome was 1. Depending on the mutation type, if it was m1, the value at that locus was 1; if it was m2, the value at that locus became -1. Each strain was assigned a score, termed a quantitative trait value, based on the difference in the number of "m1" and "m2" (score = ∑ m1 − ∑ m2), and fitness was determined by a normalized power function of the deviation from the mean score (*fitness* = 2.0 + 1.0 × (|*score* − μ|^5^)/(*max*(|*score* − μ|^5^) + 10^−6^), where *μ* is the mean score across all individuals.). Based on the trait value–derived fitness (ranging from 2.0 to 3.0), individuals with more extreme trait values had higher fitness. The number of offspring produced by each individual followed a Poisson distribution with λ = fitness, such that individuals with extreme trait values tended to produce more offspring on average. This setting imposed a diversifying selection pressure, favoring individuals at the tails of the trait value distribution.

### Refined selective model

We refined the selective model by introducing locus-specific selection coefficients. We created a dataset following a Gamma distribution to serve as the selection coefficients, with the size of the dataset corresponding to the simulated chromosome length. Since each locus had a specific selection coefficient, the assignment of values for each simulated strain was modified accordingly. The quantitative trait value was calculated as follows:

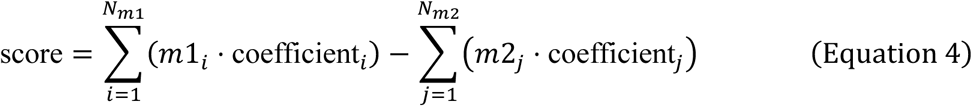

In this formula, *Nm1* and *Nm2* represent the number of m1 and m2 mutations in the simulation, respectively. This calculation differs from the previous simulations, where the trait value was simply the difference between the numbers of "m1" and "m2" mutations. We first used the previous parameters (*e.g.*, Chromosome = 1 Mb, HGT = 20 kb, and population size = 10,000) and tested three sets of experiments with a Gamma-distributed dataset, where the mean value was set to 1. The scale parameters for these experiments were set to 1×10^2^, 1×10^4^, 1×10^5^, and 1×10^6^. The timing of selection and the fitness function were the same as those used in the third simulation scenario.

To better approximate the real KP genome, we adjusted the chromosome length to 5 Mb, set the mutation rate to 10^-6^, and increased the number of HGT events to 5,, with each event transferring 20 kb, resulting in a total of 100 kb HGT per generation between each bacterium and others in the population. Each locus on the chromosome was assigned a selection coefficient drawn from a Gamma distribution, which included five million values, one for each locus on the chromosome, regardless of whether a mutation occurred at that locus. The scale for the Gamma distribution was set to 5×10^6^ for this simulation. Similarly, the mean value of the Gamma distribution was set to 1. Fitness was determined by a normalized power function of the deviation from the mean score ( *Fitness* = 2.0 + 0.5 × (|*score* − *μ*|^10^)/(*max*(|*score* − *μ*|^10^) + 10^−6^), where *μ* is the mean score across all individuals.). Based on the trait value–derived fitness (ranging from 2.0 to 2.5), individuals with more extreme trait values had higher fitness.

All the simulation scripts can be found at https://github.com/sql647/KpBACKBONE.

## Acknowledgements

This work is supported by the National Natural Science Foundation of China (NSFC) grant 32350710791 awarded to D.F. We thank many colleagues for feedback on a previous version of the manuscript.

## Supplementary Table

**Supplementary Table 1.** Enrichment of KP1_0338 high and low gene types among different host sources based on Fisher’s exact test. For each host, a one-vs-rest Fisher’s exact test was performed to evaluate whether the gene type distribution differed significantly from that of all other sources combined.

| Source | High | Low | *p*-value |
| --- | --- | --- | --- |
| Animal | 10 | 23 | 6.907177e-01 |
| Bird | 6 | 8 | 2.196240e-01 |
| Cat | 9 | 35 | 4.892939e-01 |
| Cow | 153 | 376 | 1.523384e-01 |
| Dog | 27 | 75 | 1.000000e+00 |
| Environment | 60 | 198 | 2.272826e-01 |
| Food | 12 | 21 | 2.313088e-01 |
| Horse | 58 | 67 | 9.531088e-07 |
| Mouse | 1 | 16 | 5.529954e-02 |
| Pig | 28 | 112 | 7.438013e-02 |
| Poultry | 9 | 59 | 1.128966e-02 |
| Water | 8 | 79 | 5.870439e-05 |

## Supplementary Figures

**Supplementary Figure 1.**
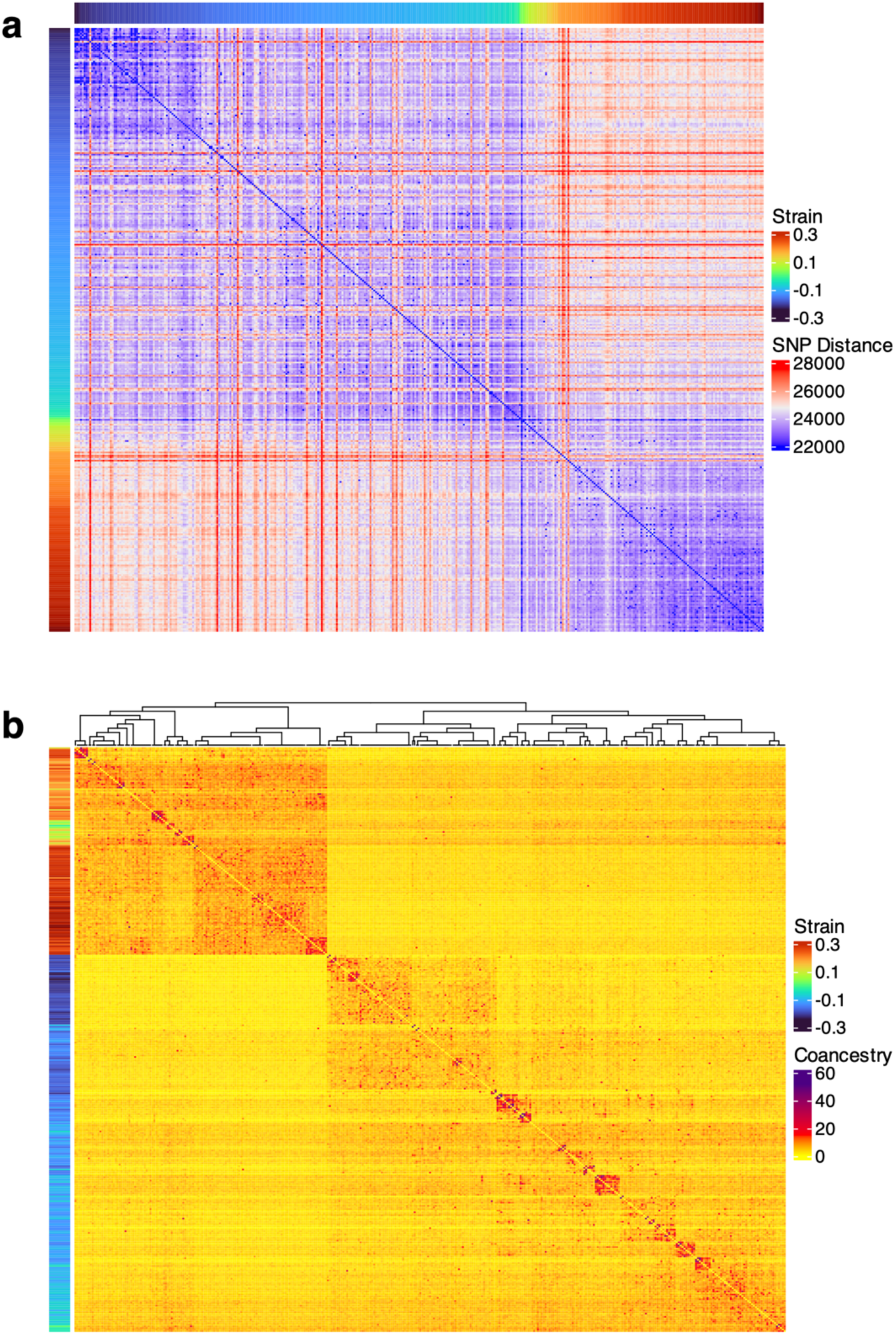
Genomic characteristics of the non-redundant KP dataset. (a) SNP distance matrix illustrating the genetic distances between strains in the non-redundant dataset, with strains ordered based on their PC1 values. The minimum SNP distance observed is 20,000 SNPs. (b) fineSTRUCTURE analysis of the non-redundant KP dataset, highlighting population structure and clustering patterns. The “strain” bar on the left represents the PC1 values of the respective strains.

**Supplementary Figure 2.**
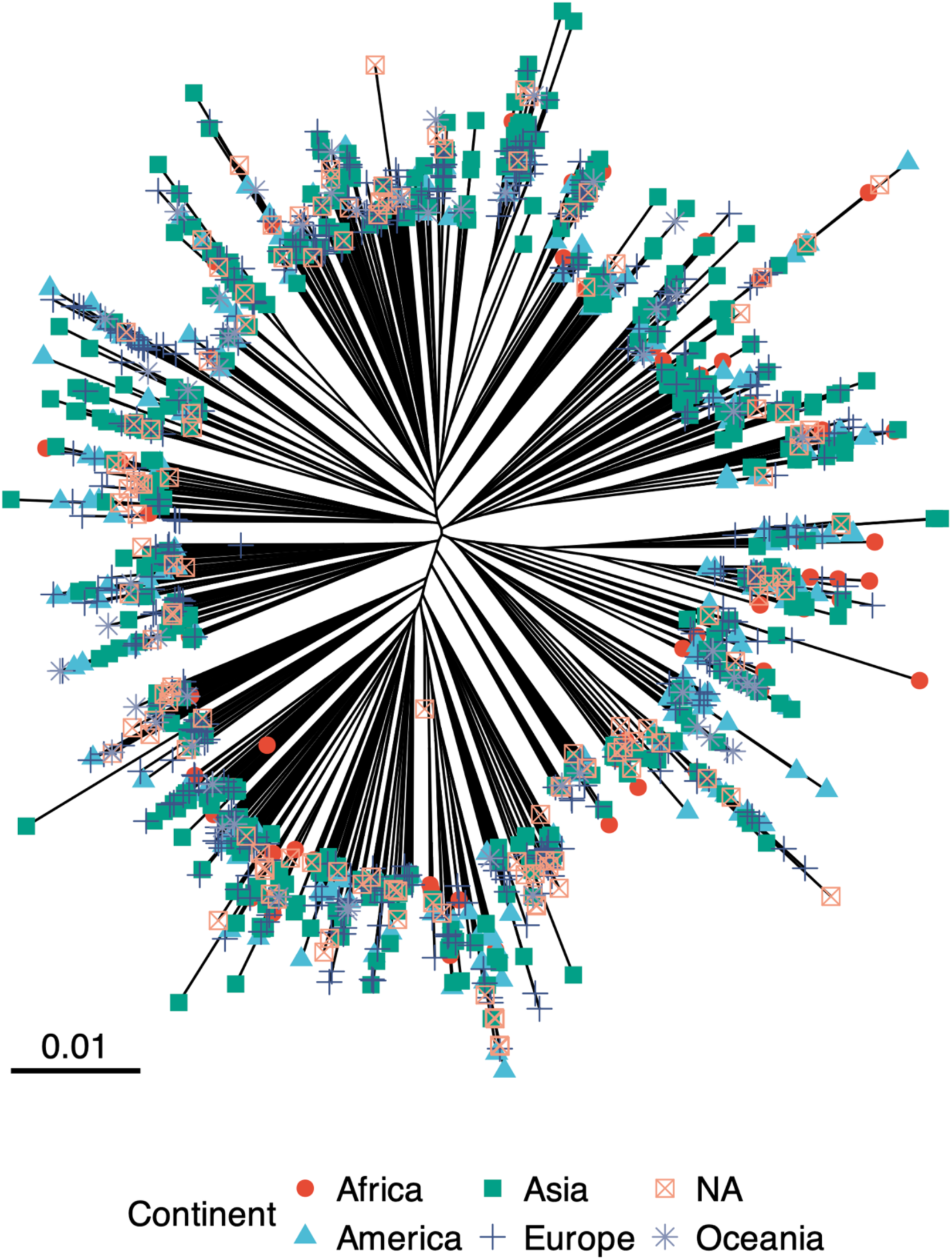
Phylogenetic tree of the complete KP dataset. A maximum-likelihood phylogenetic tree constructed based on core SNPs from the full KP dataset, with branches color-coded by continental origin to illustrate the global distribution of isolates. NA: no information available.

**Supplementary Figure 3.**
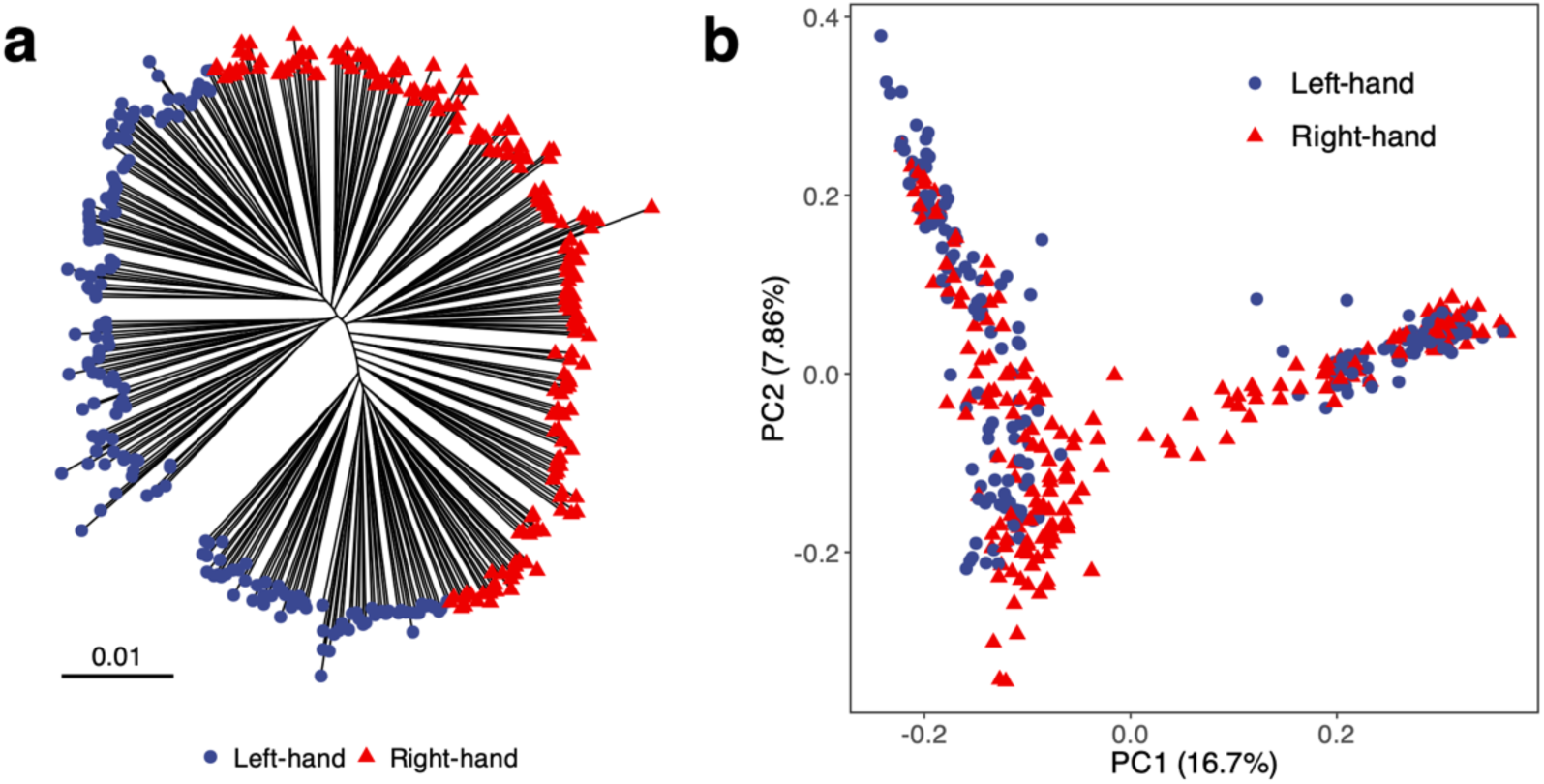
(a) Phylogenetic tree of the non-redundant KP dataset divided into two subsets based on the backbone. (b) PCA of the non-redundant KP dataset colored according to two sub-groups.

**Supplementary Figure 4.**
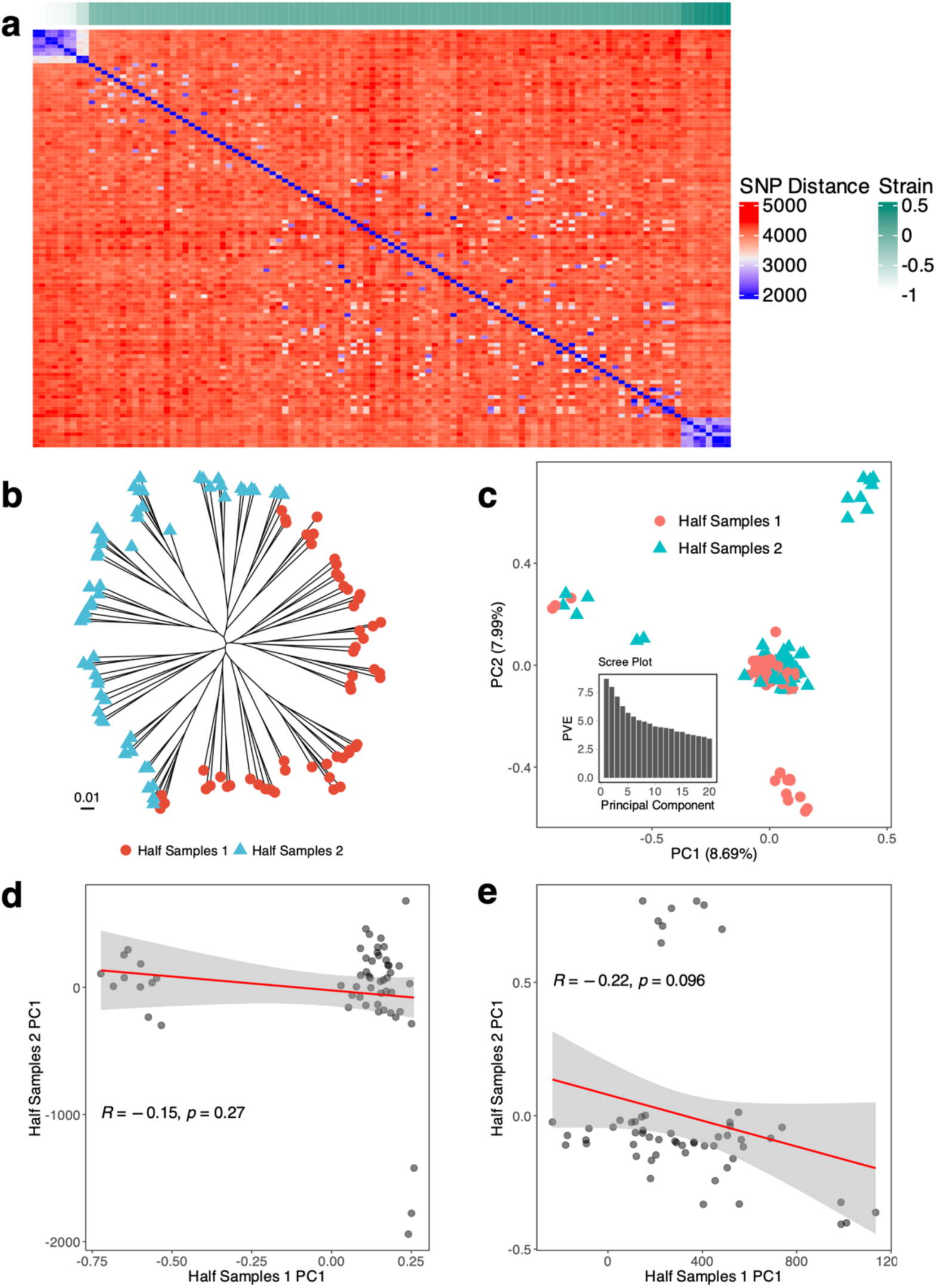
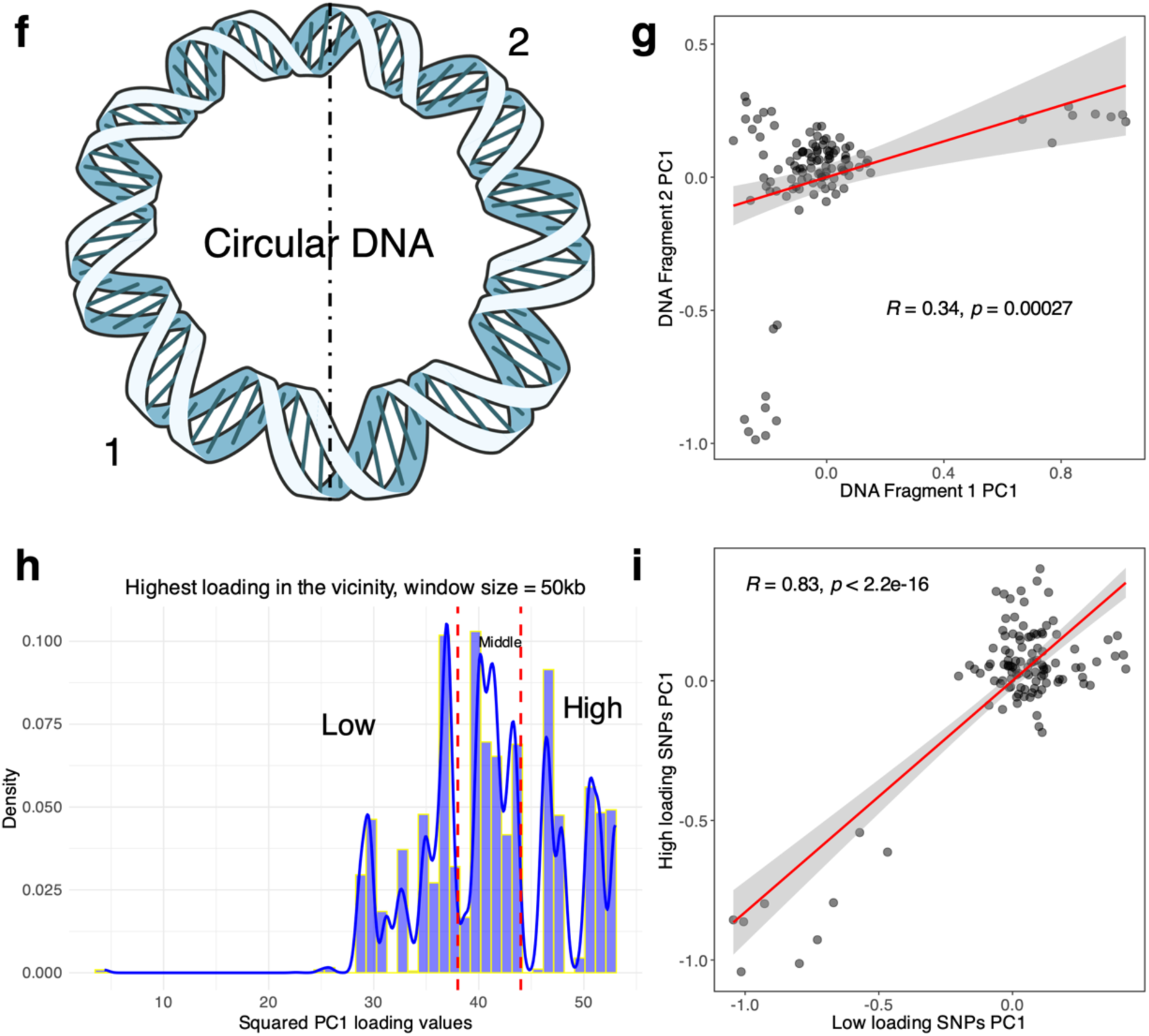
Three types of half-matching tests for the clonal simulation. (a) SNP distance matrix for the clonal simulation. The minimum SNP distance observed is 2,000 SNPs. (b–e) Half-strain matching: (b) Phylogenetic tree of the non-redundant clonal simulation divided into 2 subsets. (c) PCA of the non-redundant clonal simulation data. (d) Correlation of PC1 from group 2 projected onto group 1 results. (e) Correlation of PC1 from group 1 projected onto group 2 results. (f–g) Half-genome matching: (f) Division of the genome into two fragments (Fragment 1 and Fragment 2). (g) Correlation of PC1 between the two fragments. (h–i) Partial-variants matching: (h) Histogram of SNP loadings on PC1, dividing SNPs into high- and low-loading groups. (i) Correlation of PC1 between high- and low-loading SNP groups.

**Supplementary Figure 5.**
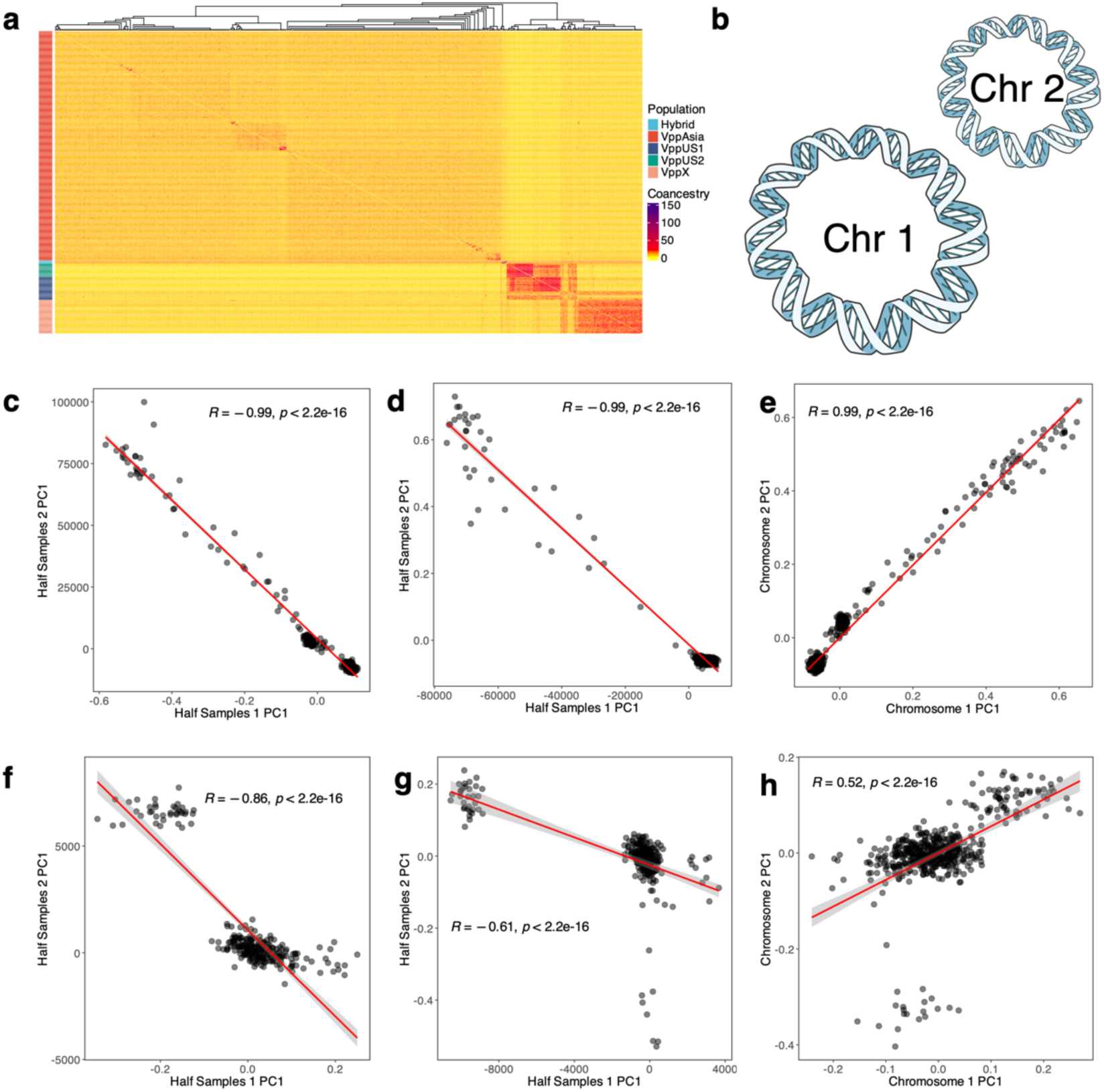
Half-matching tests for VP datasets. (a) fineSTRUCTURE analysis assigning populations for the non-redundant VP dataset. (b) Representation of VP Chromosome 1 and Chromosome 2, with proportions reflecting their actual sizes. (c-e) Half-strain matching for the non-redundant VP dataset: (c) Correlation of PC1 from group 2 projected onto group 1 results; (d) Correlation of PC1 from group 1 projected onto group 2 results. Half-chromosome matching for the non-redundant VP dataset: (e) Correlation of PC1 between the two chromosomes. (f-h) Half-matching for VppAsia dataset: (f) Correlation of PC1 from group 2 projected onto group 1 results; (g) Correlation of PC1 from group 1 projected onto group 2 results. Half-chromosome matching for VppAsia dataset: (h) Correlation of PC1 between the two chromosomes.

**Supplementary Figure 6.**
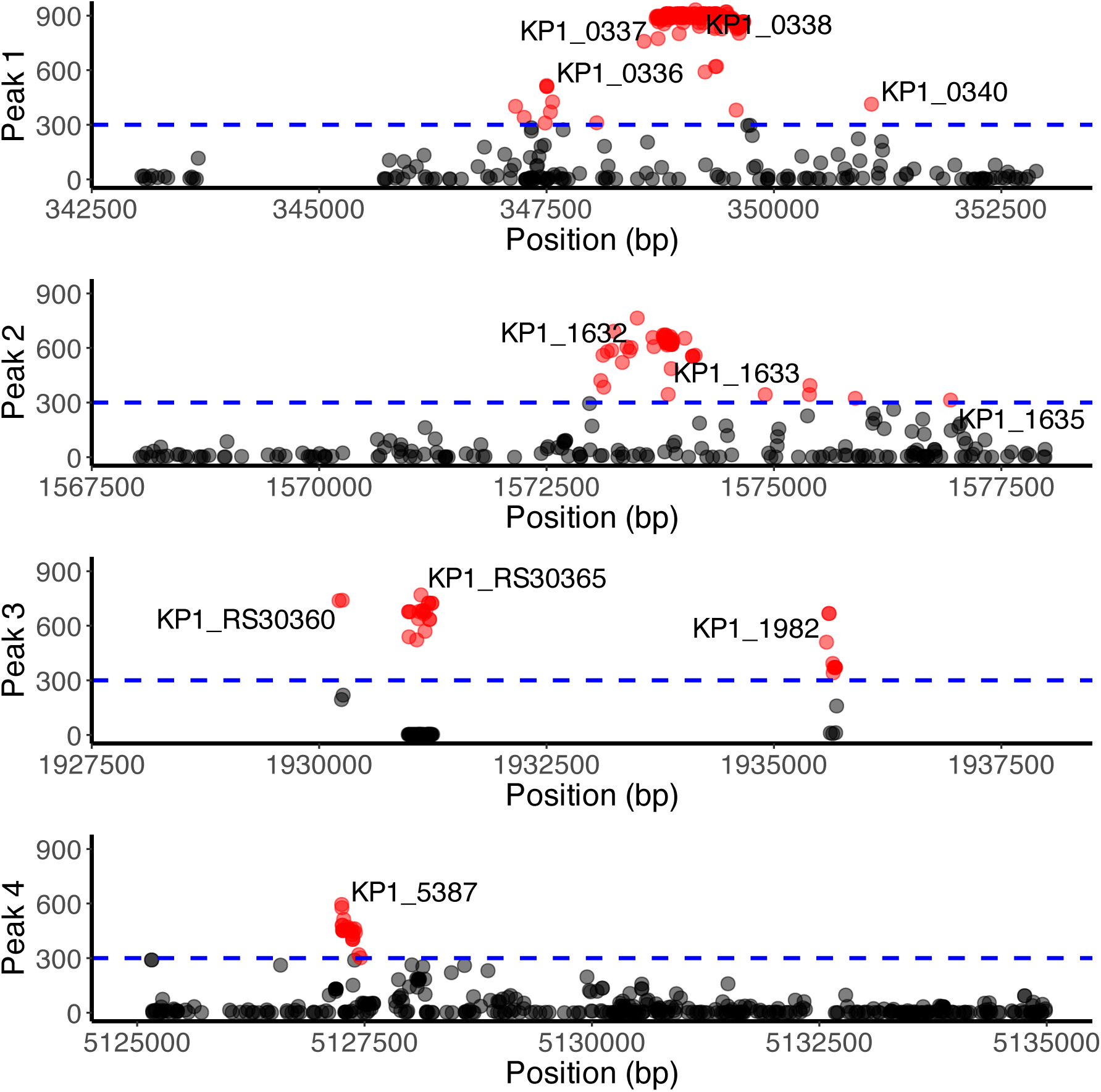
Zoomed-in views of four high-loading SNP peaks identified across the genome.

**Supplementary Figure 7.**
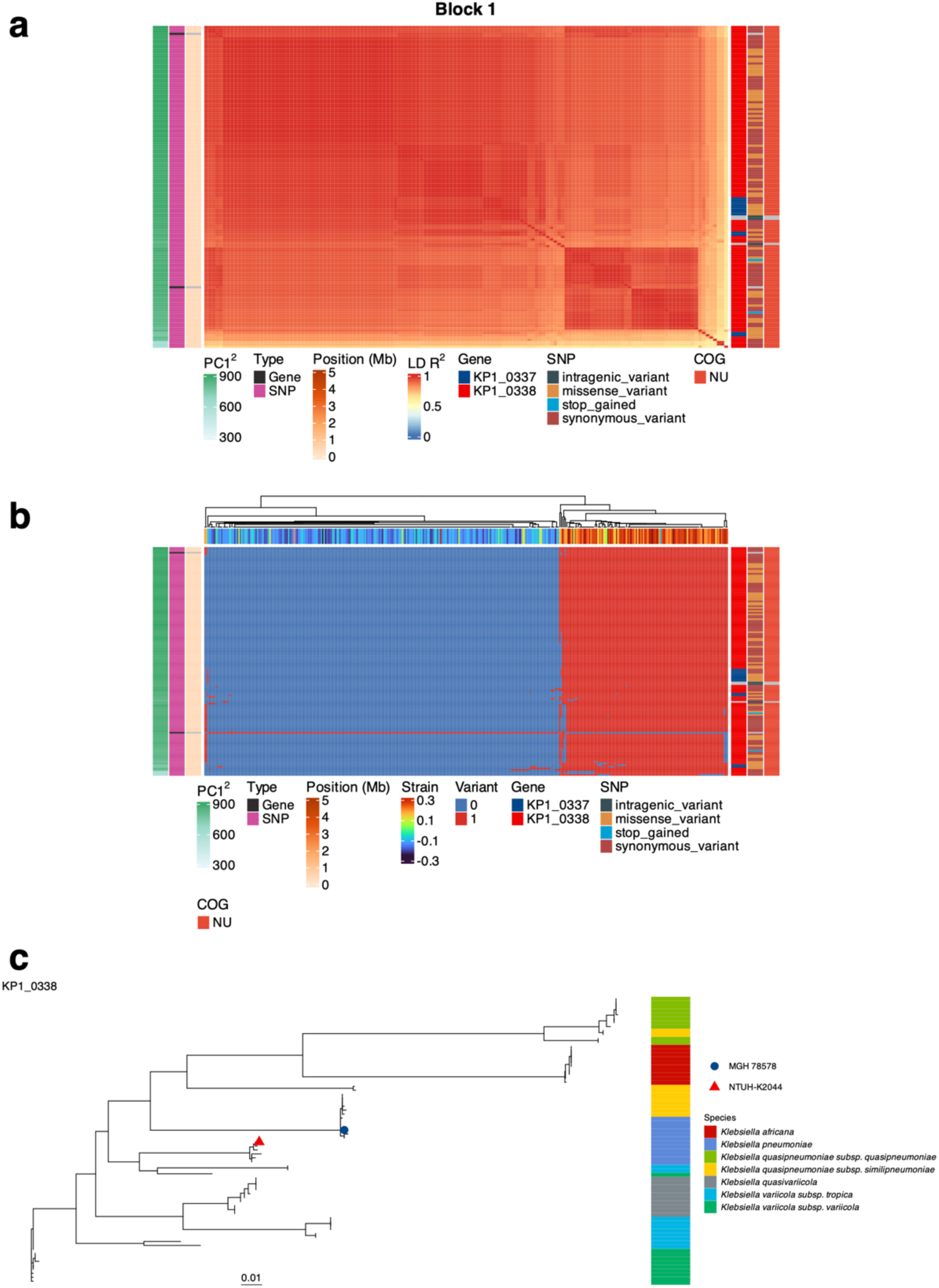
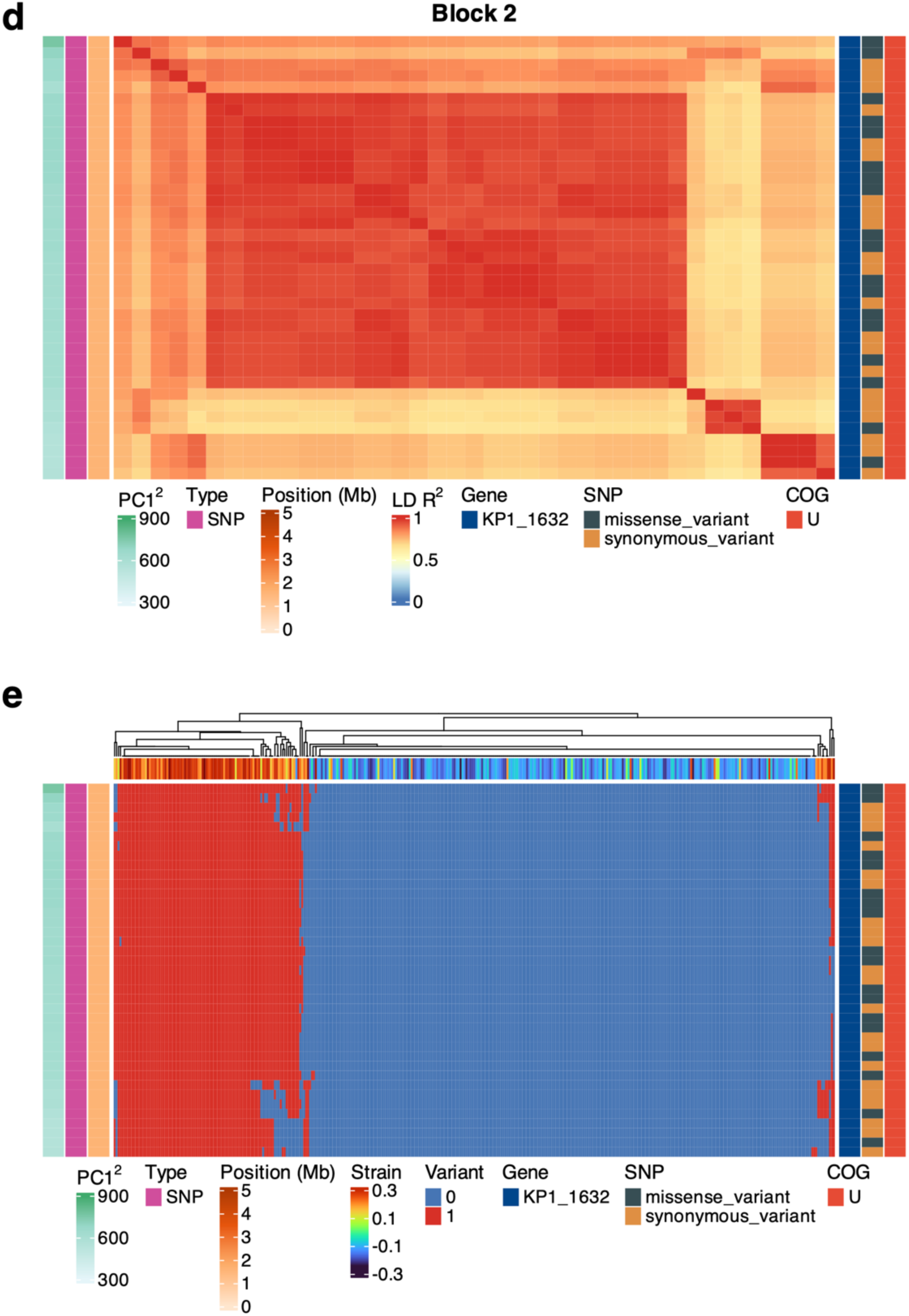
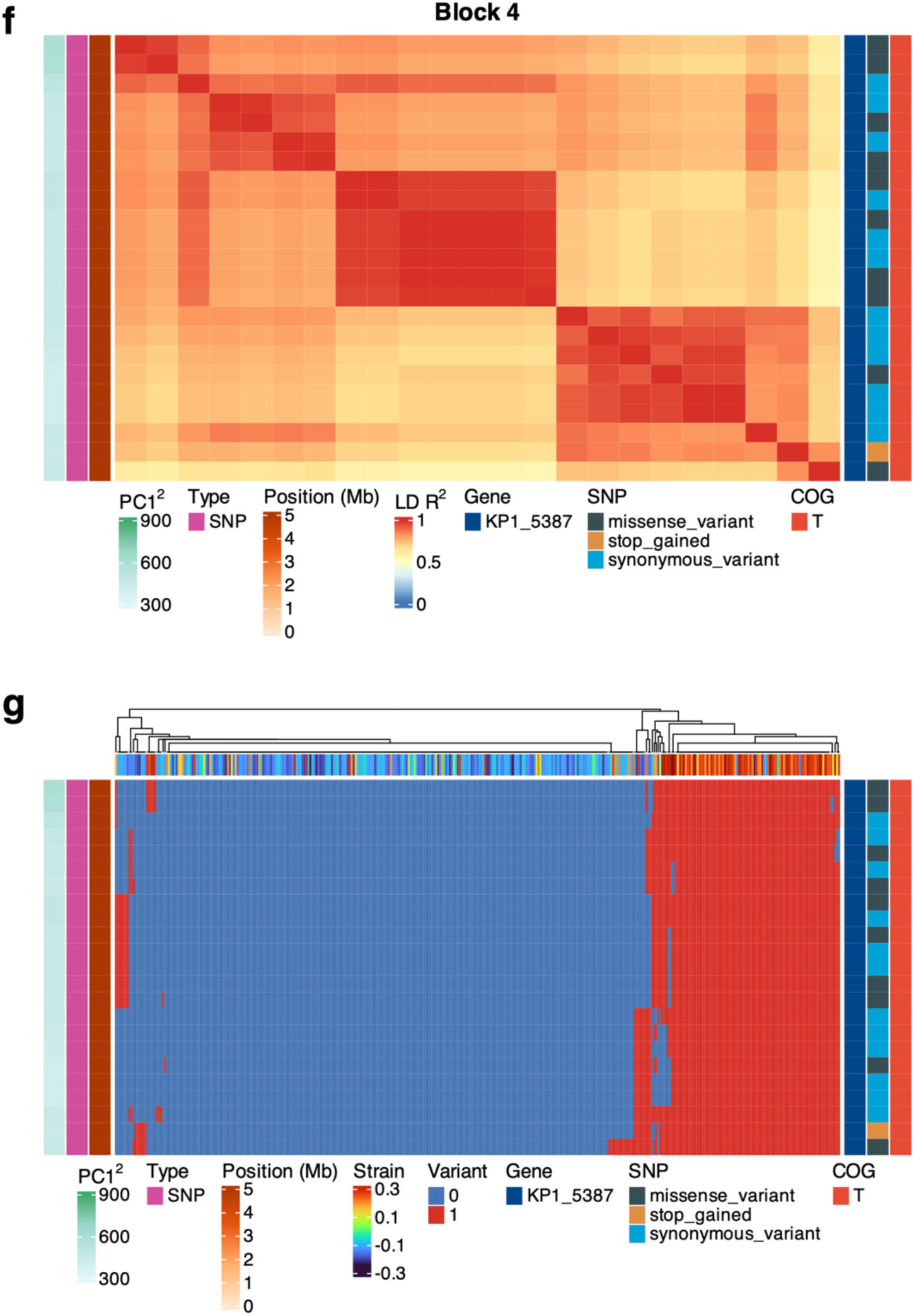
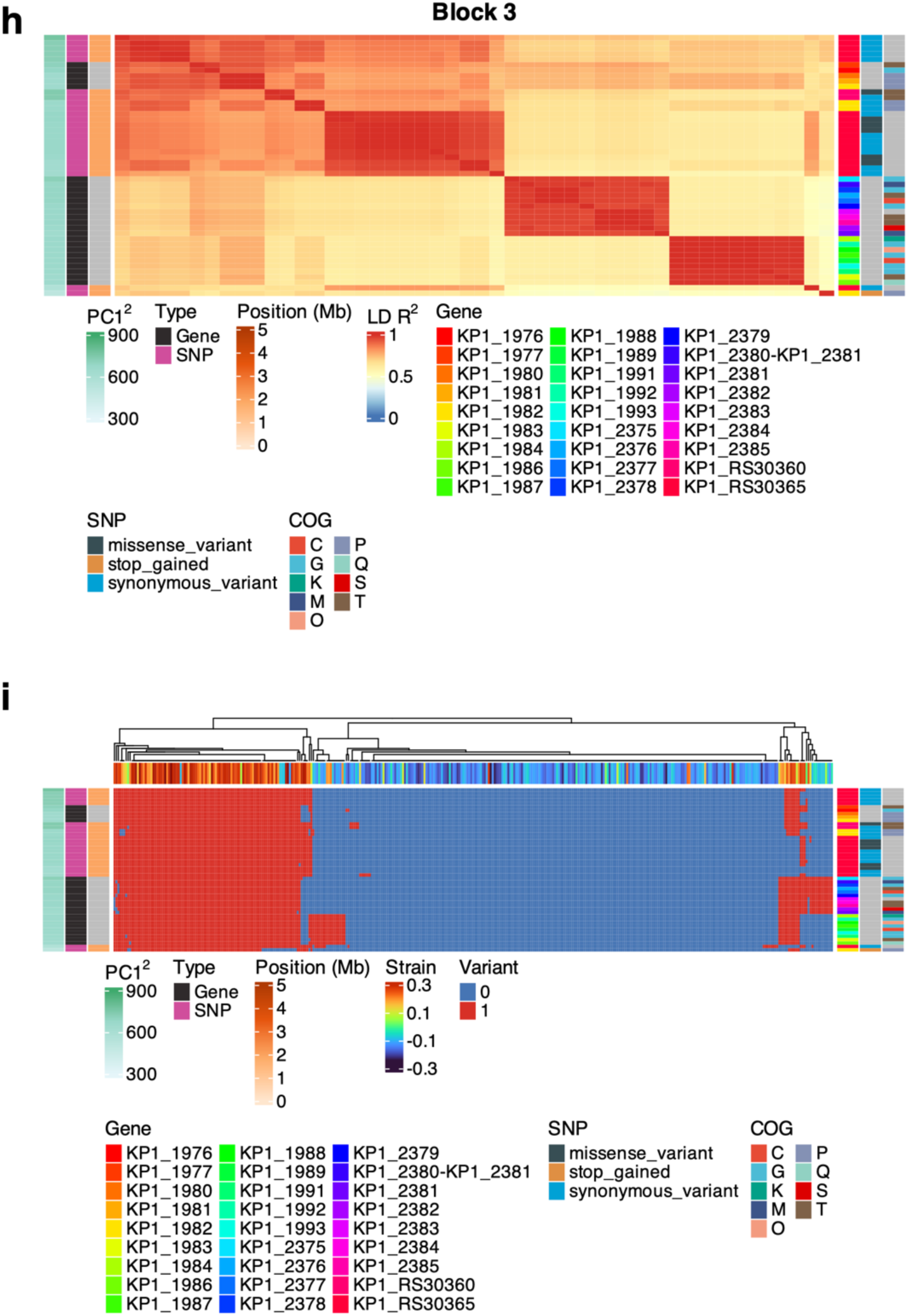
Overview of blocks 1–4 from LD analysis with variant status heatmaps. Block 1 (a-b): (a) SNPs in KP1_0337, KP1_0338, and associated accessory genes form a high LD block. (b) Variant (1/0) heatmap shows clear differentiation in gene presence/absence or SNP minor/major alleles across isolates. (c) Multi-species phylogenetic tree of the KP1_0338 gene. The annotation strip indicates the species of each strain. Block 2 (d-e): (d) High LD observed for SNPs in KP1_1632. (e) Variant (1/0) heatmap highlights distinct patterns of gene presence/absence or SNP minor/major alleles across isolates. Block 4 (h-i): (h) High LD observed for variants in KP1_5387, forming a small, highly linked cluster. (i) Variant (1/0) heatmap shows distinct variation in gene presence/absence or SNP minor/major alleles across isolates. Block 3 (f-g): (f) SNPs in KP1_RS09345, KP1_RS30360, KP1_1982, and accessory genes form tightly linked clusters. (g) Variant (1/0) heatmap reveals clear differentiation in gene presence/absence or SNP minor/major alleles across isolates.

**Supplementary Figure 8.**
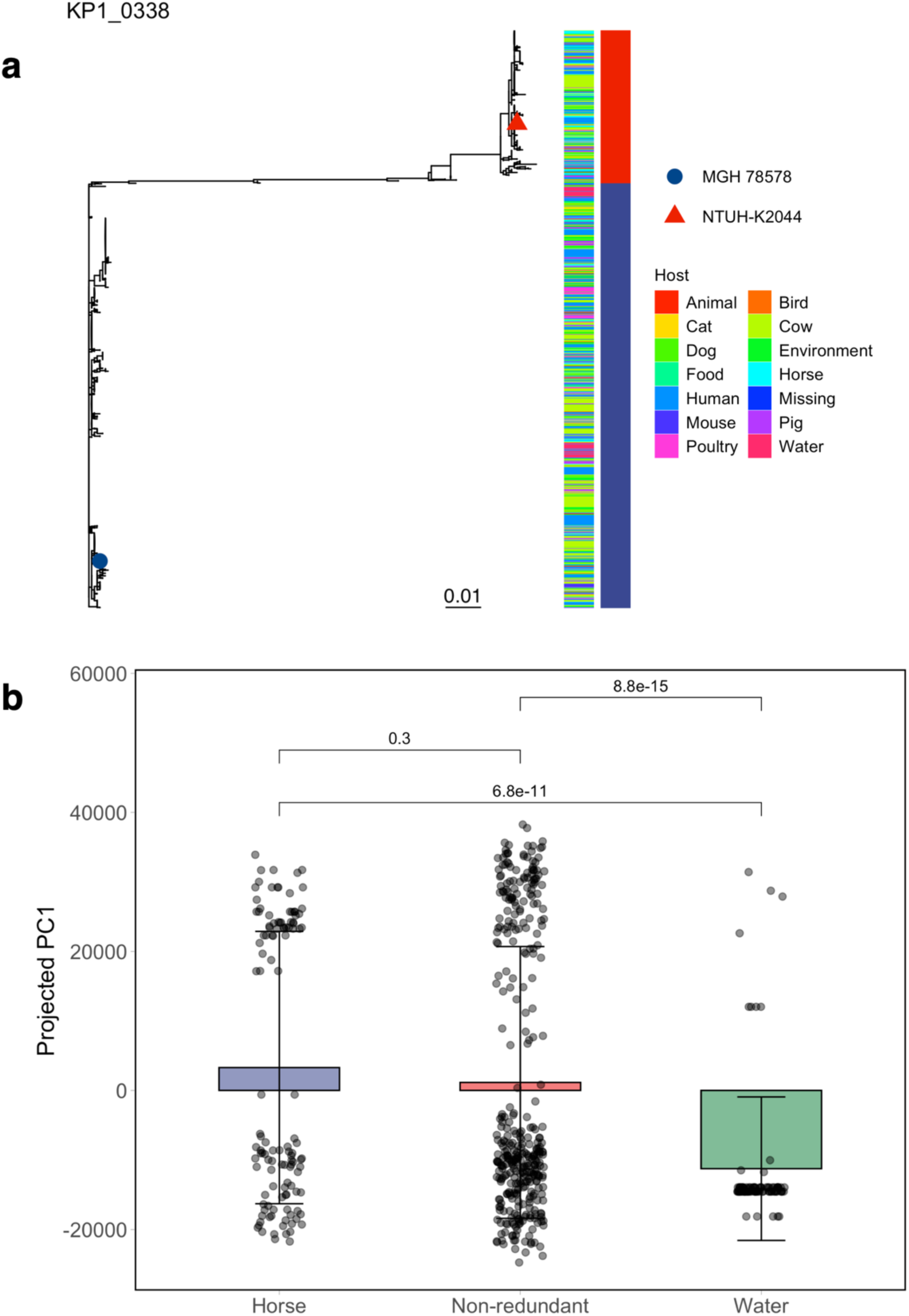
Phylogenetic tree of KP1_0338 within KP and host origin profiles. (a) Phylogenetic tree of the KP1_0338 gene based on sequences from the multi-host KP dataset. Two annotation strips indicate host origin and KP1_0338 classification (High vs. Low) based on phylogenetic clustering. (b) Projected the multi-host dataset onto the non-redundant dataset. Each bar represents the mean PC1 value for isolates from a given host, with error bars indicating the standard deviation.

**Supplementary Figure 9.**
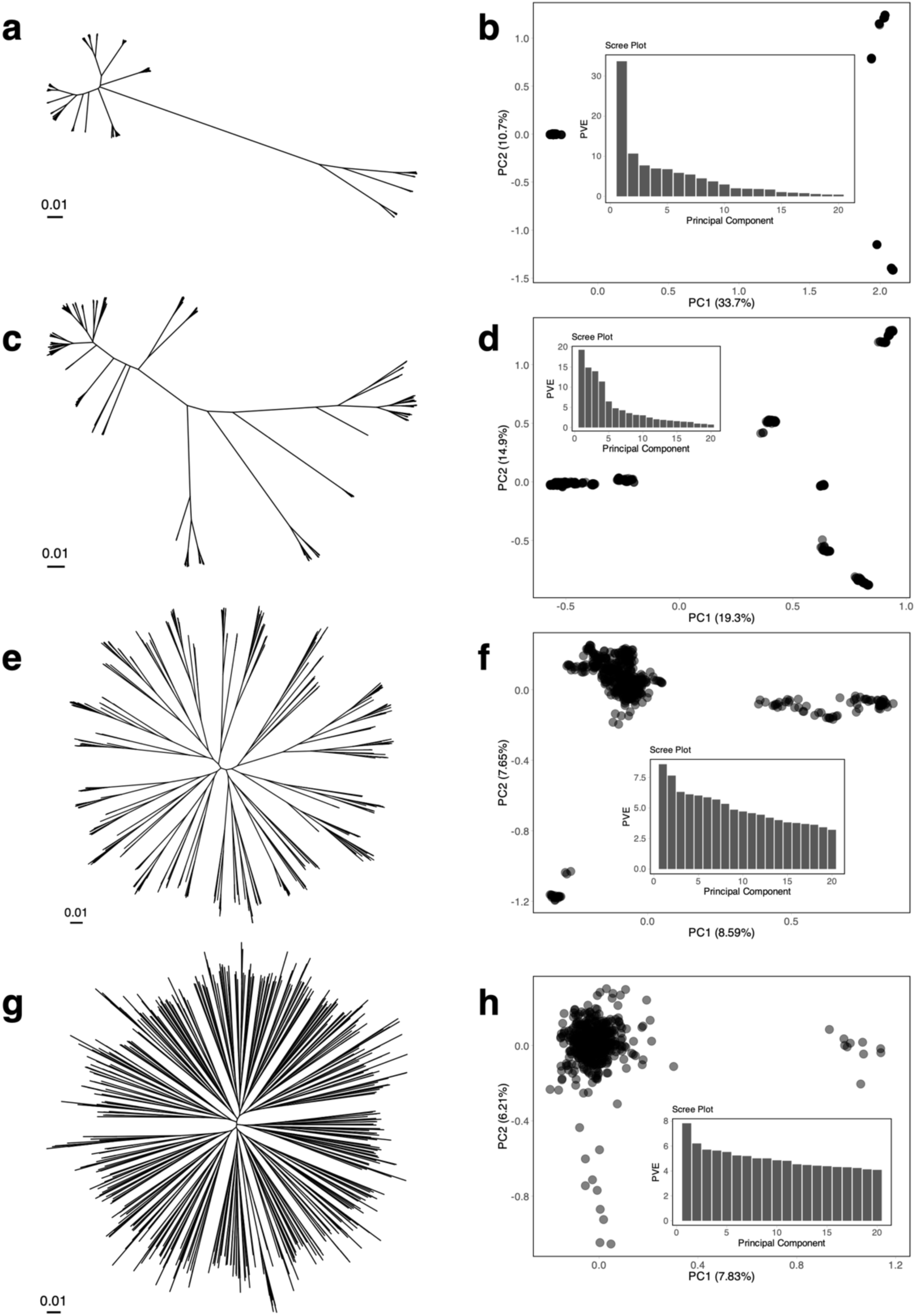
Clonal haploid bacteria model with varying HGT lengths and a population size of 2000. Phylogenetic trees, PCA plots, and scree plots for a population of 2000 individuals under different HGT lengths. Panels (a-h) show results for: (a-b) 100 bp, (c-d) 1 kb, (e-f) 10 kb, (g-h) 100 kb.

**Supplementary Figure 10.**
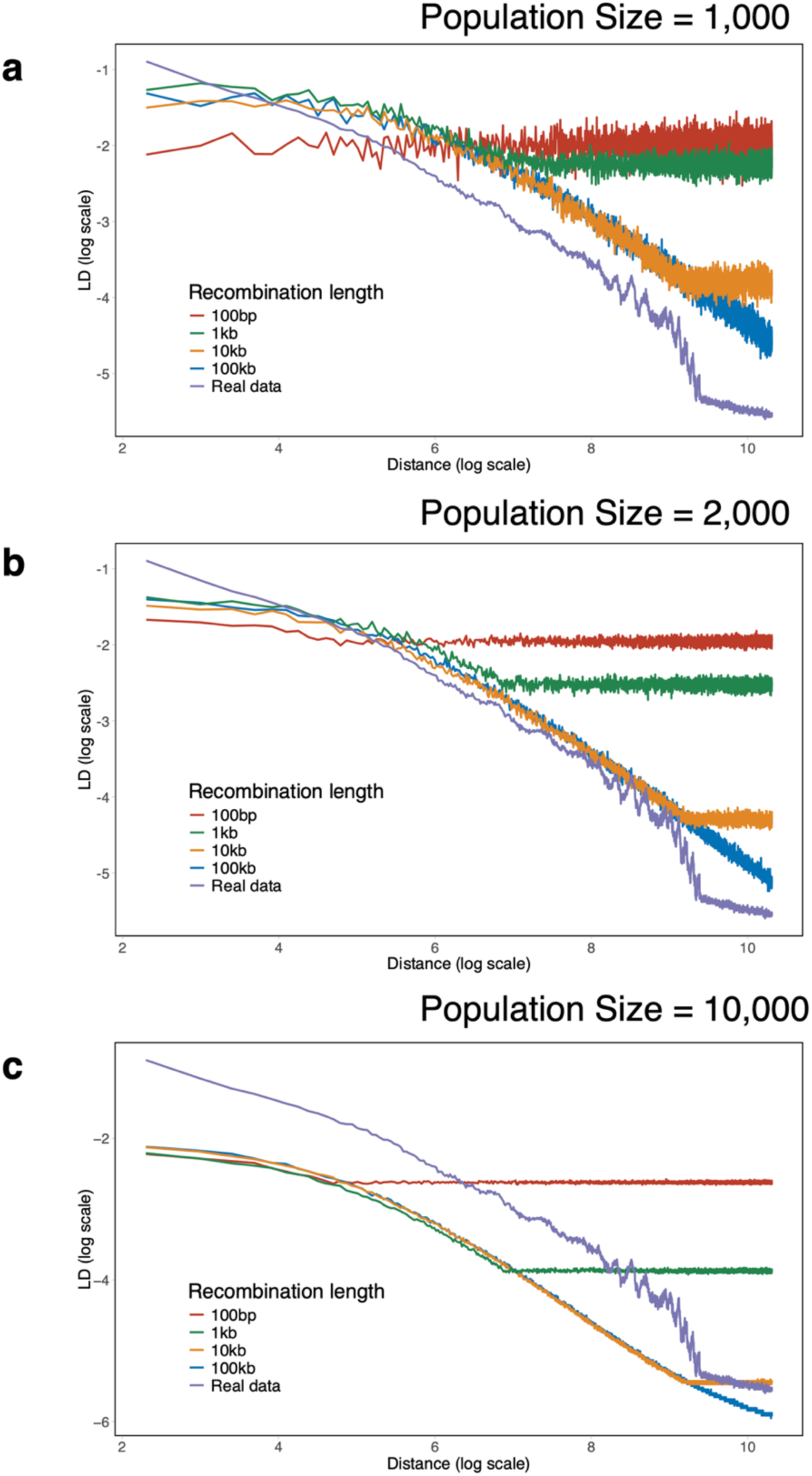
LD decay under different HGT lengths in clonal haploid bacteria models. (a-c) Comparison of LD decay between different HGT lengths (100 bp, 1 kb, 10 kb, and 100 kb) and the non-redundant KP dataset. (a) Population size of 1,000. (b) Population size of 2,000. (c) Population size of 10,000. The x-axis represents the SNP distance (log scale), and the y-axis represents the LD value (*R^2^*) on a log scale.

**Supplementary Figure 11.**
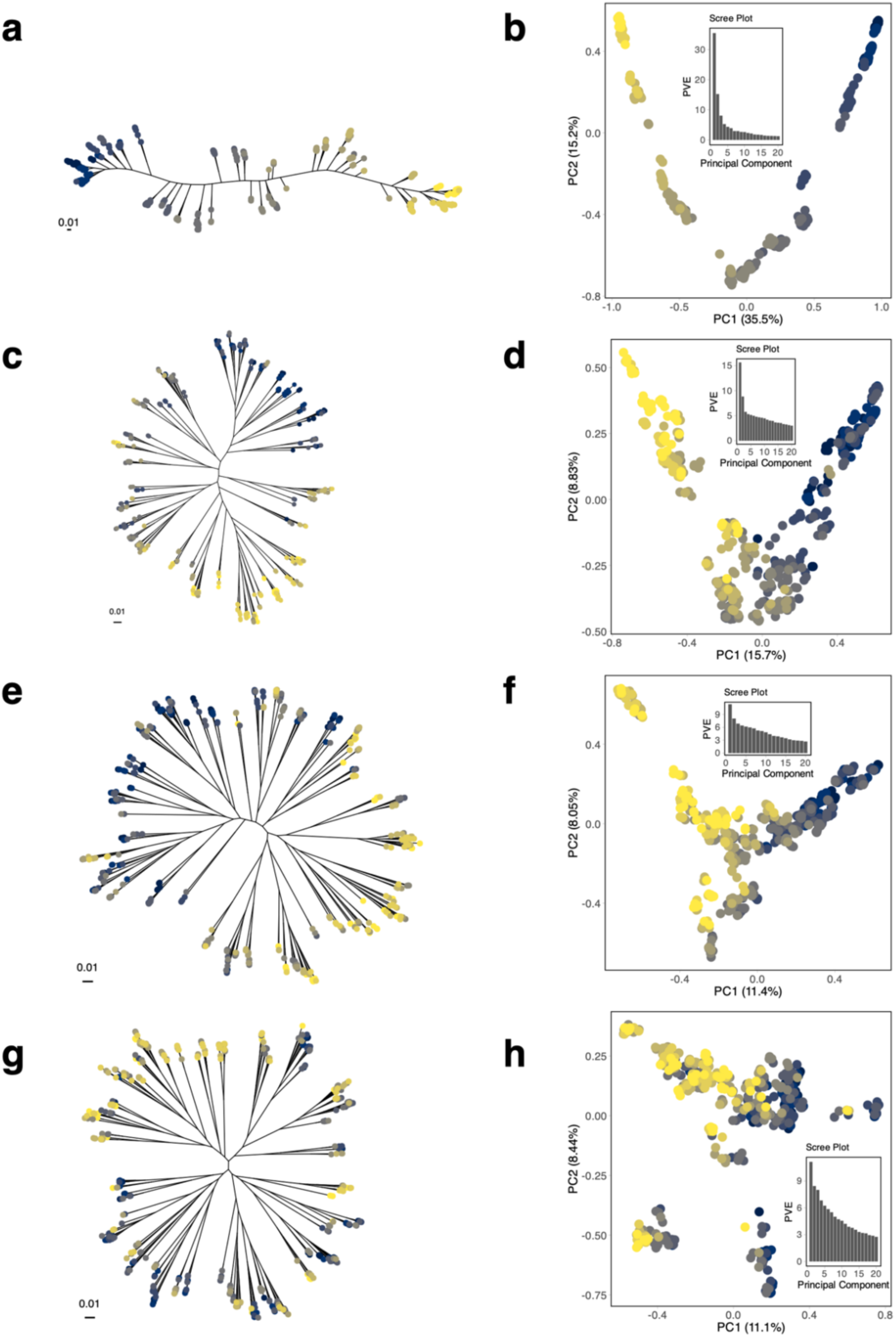
Linear stepping-stone model with varying numbers of migrants and a population size of 2000. Phylogenetic trees, PCA plots, and scree plots for a model with HGT of 10 kb. The population consists of 10 subpopulations (200 individuals each). Panels (a–h) show results for different numbers of migrants exchanged between neighbor subpopulations: (a-b) 1 migrant, (c-d) 10 migrants, (e-f) 20 migrants, (g-h) 50 migrants.

**Supplementary Figure 12.**
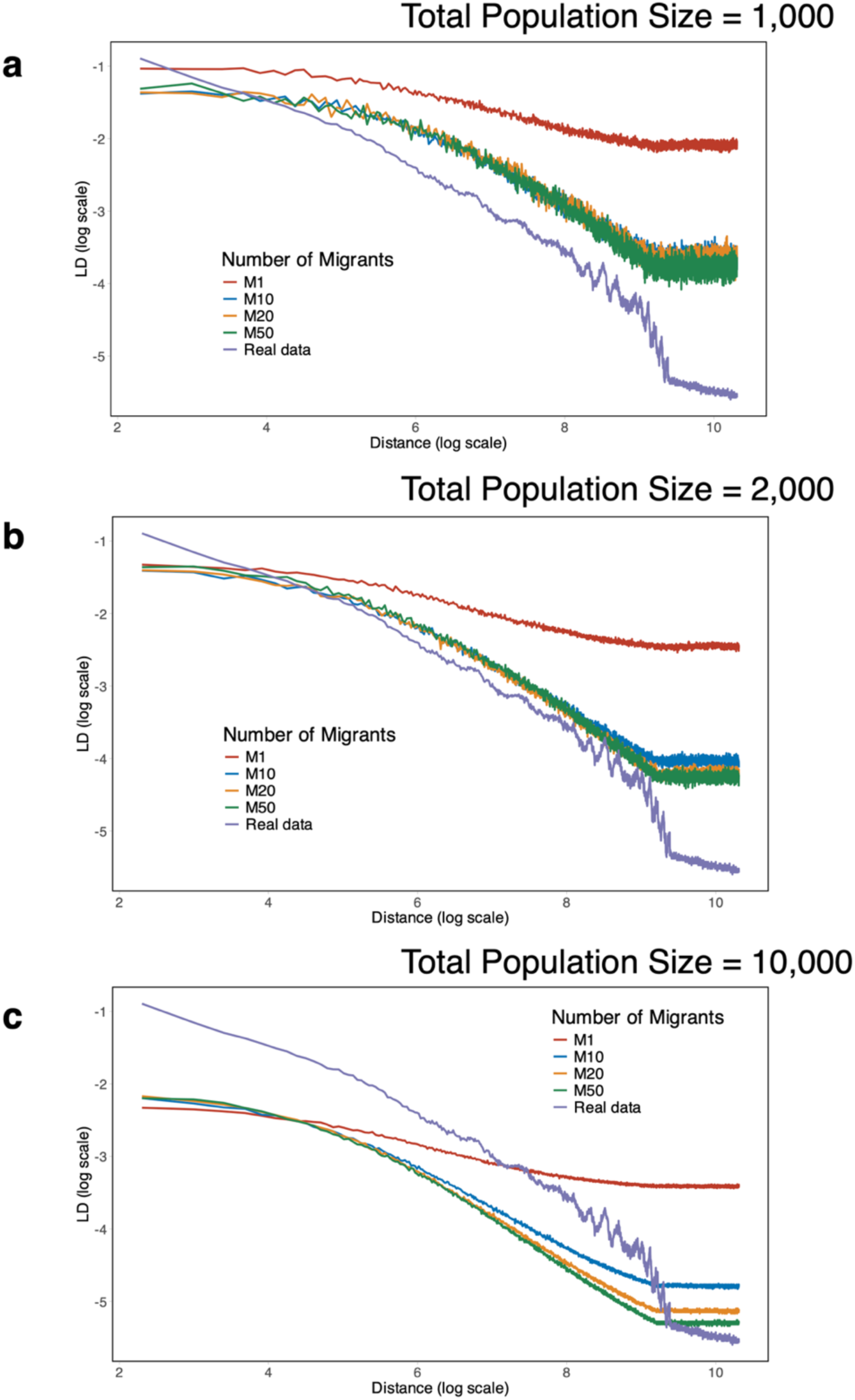
LD decay profiles are compared between linear stepping-stone models with varying numbers of migrants (1, 10, 20, 50) and the non-redundant KP dataset. Panels illustrate results for different total population sizes: (a) 2000 individuals, (b) 10,000 individuals, (c) 100,000 individuals. Each model includes HGT of 10 kb. The x-axis represents the SNP distance (log scale), and the y-axis represents the LD value (*R^2^*) on a log scale.

